# Landscape of multi-nucleotide variants in 125,748 human exomes and 15,708 genomes

**DOI:** 10.1101/573378

**Authors:** Qingbo Wang, Emma Pierce-Hoffman, Beryl B. Cummings, Konrad J. Karczewski, Jessica Alföldi, Laurent C. Francioli, Laura D. Gauthier, Andrew J. Hill, Anne H. O’Donnell-Luria, Genome Aggregation Database (gnomAD) Production Team, Genome Aggregation Database (gnomAD) Consortium, Daniel G. MacArthur

## Abstract

Multi-nucleotide variants (MNVs), defined as two or more nearby variants existing on the same haplotype in an individual, are a clinically and biologically important class of genetic variation. However, existing tools for variant interpretation typically do not accurately classify MNVs, and understanding of their mutational origins remains limited. Here, we systematically survey MNVs in 125,748 whole exomes and 15,708 whole genomes from the Genome Aggregation Database (gnomAD). We identify 1,996,125 MNVs across the genome with constituent variants falling within 2 bp distance of one another, of which 31,510 exist within the same codon, including 405 predicted to result in gain of a nonsense mutation, 1,818 predicted to rescue a nonsense mutation event that would otherwise be caused by one of the constituent variants, and 16,481 additional variants predicted to alter protein sequences. We show that the distribution of MNVs is highly non-uniform across the genome, and that this non-uniformity can be largely explained by a variety of known mutational mechanisms, such as CpG deamination, replication error by polymerase zeta, or polymerase slippage at repeat junctions. We also provide an estimate of the dinucleotide mutation rate caused by polymerase zeta. Finally, we show that differential CpG methylation drives MNV differences across functional categories. Our results demonstrate the importance of incorporating haplotype-aware annotation for accurate functional interpretation of genetic variation, and refine our understanding of genome-wide mutational mechanisms of MNVs.

## Introduction

Multi-nucleotide variants (MNVs) are defined as clusters of two or more nearby variants existing on the same haplotype in an individual^1,2^ (Fig. 1a). When variants in an MNV are found within the same codon, the overall impact may differ from the functional consequences of the individual variants^3^. For instance, the two variants depicted in Fig. 1b are each predicted individually to have missense consequences, but in combination result in a nonsense variant. Such cases, which would be missed by virtually all existing tools for clinical variant annotation, can result both in missed diagnoses and false positive pathogenic candidates in analyses of families affected by genetic diseases^1,2^.

**Figure 1.**
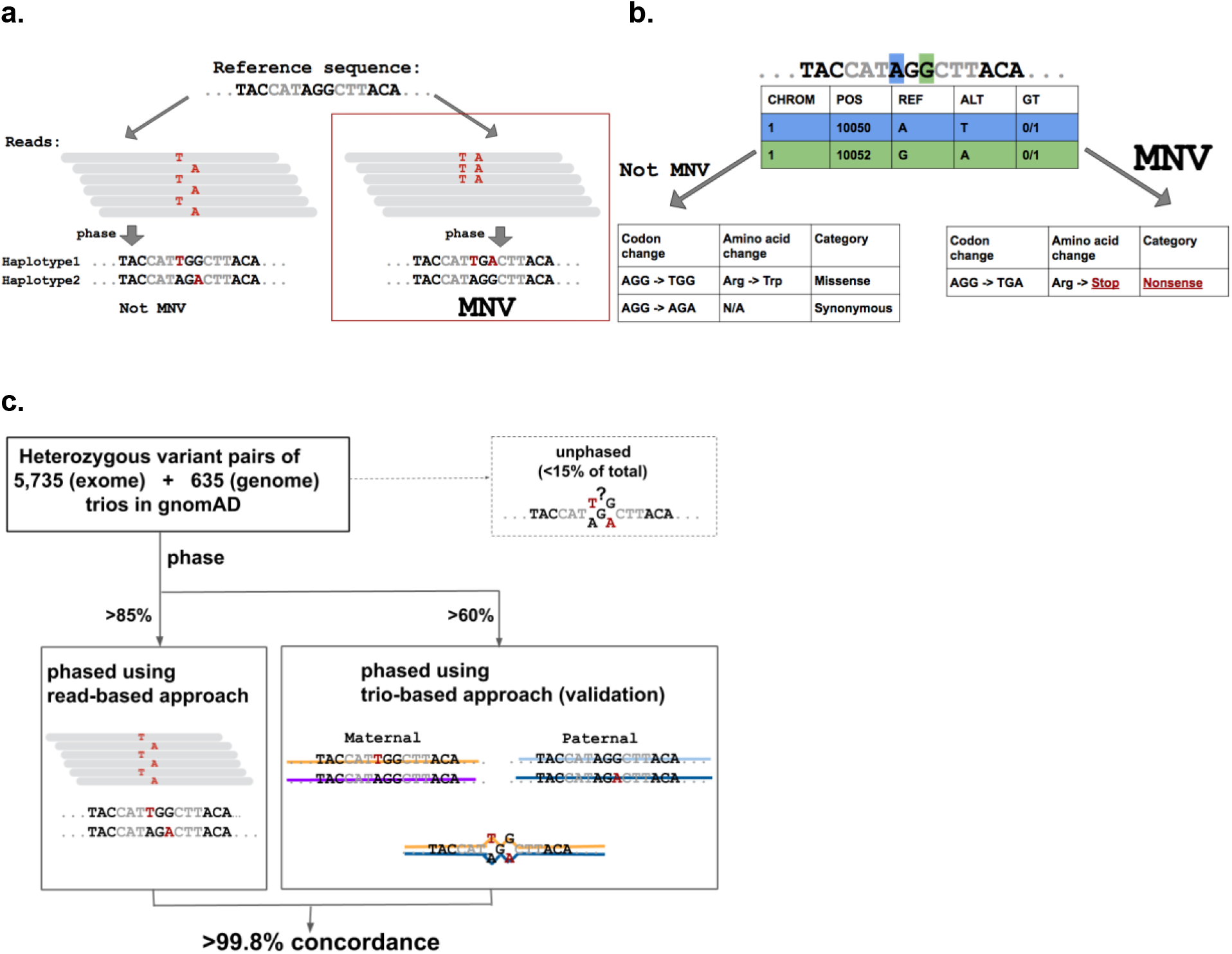
Definition and an example of MNVs, and validation of phasing sensitivity. **a**, Definition and an example of an MNV. In this manuscript, an MNV is defined as two or more nearby variants existing on the same haplotype in the same individual. **b**, Impact of MNVs in coding regions. The amino acid change caused by an MNV can be different from either of the individual single nucleotide variants, which creates the potential for missannotation of the functional consequence of variants. **c**, Graphical overview of the analysis of phasing sensitivity and specificity using trio samples from our gnomAD callset. We identified all heterozygous variant pairs that pass quality control (Methods) and compared the phase information assigned by read based phasing with that of trio based phasing.

MNV identification tools^4–8^ have been applied to databases of human genetic variation at varying scales, including 1000 Genomes^9^ Phase 3 (2,504 individuals with high coverage exome and low coverage genome sequencing data), and the Exome Aggregation Consortium^1^ (60,706 individuals with high coverage exome data). Together, these analyses identified over 10,000 MNVs altering protein sequences, demonstrating the pervasive nature of MNV annotation in population level data. Additionally, analysis of the 1000 Genomes dataset highlighted differences in the frequencies of MNVs depending on sequence context^10^. In combination with yeast experiments^11–13^, biological mechanisms that account for the enrichment of specific types of MNVs, such as DNA replication error by polymerase zeta, have been suggested.

Studies of newly occurring *(de novo)* MNVs have also been performed using trio datasets^2,14–16^; analysis of 283 trios with whole genome sequence data^16^ confirmed that MNV events occur much more frequently than expected by random chance. By focusing on non-coding regions, this study also highlighted potentially different mechanisms that dominate MNV generation depending on the genomic region and the distance between the two constitutive variants. As part of the Deciphering Developmental Disorders (DDD) study^17^, Kaplanis et. al.^2^ analyzed exome sequence data from over 6,000 trios to quantify the pathogenic impact of MNVs in developmental disorders, showing that such variants are substantially more likely to be deleterious than SNVs and further clarifying the mutational mechanisms that generate them. These analyses also have provided estimates of the germline MNV rate per generation, falling into a consistent range of 1-3% of the SNV rate. Although these studies have provided valuable information about the mutational origins and functional impact of MNVs, to date there has been no analysis that investigated MNVs across the entire genome (including non-coding regions) in many thousands of deeply sequenced individuals, limiting our understanding of the genome-wide profile and complete frequency distribution of this class of variation.

Here, we present the analysis of the largest collection of MNVs assembled to date, along with clinical interpretation of MNVs from over 6,000 sequenced individuals from rare disease families. We also provide gene-level statistics on MNVs and describe the distribution of MNVs by functional consequence and by gene-level constraint. Finally, to enhance our understanding of MNV mechanisms, we examine the distributions of MNVs stratified by more than ten different functional annotations across the human genome, as well as estimates of the genome-wide per-base frequencies of the dominant mutational processes generating MNVs.

## Results

### Read-based phasing for identification of MNVs

Identification of MNVs requires the constituent variants to be properly phased - that is, to be identified accurately as either both occurring on the same haplotype (in *cis*) or on two different haplotypes (in *trans*). Phasing can be performed following three broad strategies: read-based phasing^18^, which assesses whether nearby variants co-segregate on the same reads in DNA sequencing data; family-based phasing^19^, which assesses whether pairs of variants are co-inherited within families; and population-based phasing^20^, which leverages haplotype sharing between members of a large genotyped population to make a statistical inference of phase. Read-based phasing is particularly effective for pairs of nearby variants, making it suitable for the analysis of MNVs.

For this project, we generated read-based phasing results for variants in the Genome Aggregation Database (gnomAD) v2.1 callset using GATK HaplotypeCaller^21^, yielding 125,748 human whole exomes and 15,708 genomes with local phase information; the properties of this callset are described in detail in an accompanying manuscript^22^. To assess phasing accuracy, we used 5,785 family trios with exome sequencing data and 635 family trios with whole genome sequencing data that largely overlapped with the gnomAD 2.1 release data. We calculated the phasing sensitivity, defined as the fraction of heterozygous variant pairs that have read-based phase information assigned for both variants, and found that it was 85.2% for adjacent heterozygous variant pairs, reflecting the stringent haplotype-calling criteria of GATK^21^ (Supplementary Table 1). We used Phase-By-Transmission (PBT)^19^, a family-based phasing method (Fig. 1c), to assess our phasing specificity, and found that over 99.8% of the MNVs identified with read-based phasing were consistent with the PBT trio-based phasing. The sensitivity and specificity of our read-based phasing remained high even when the two variants of the MNV were 10 bp apart (80.6% and 99.4%; Supplementary Fig. 1 and Supplementary Table 1). These results demonstrate high specificity and sensitivity for the detection of MNV events across the genome.

### Functional impact of MNVs

In order to provide an overview of the functional impact of MNVs (Fig. 1b), we examined all phased high-quality SNV pairs (i.e. SNV pairs that pass stringent filtering criteria; see Methods) within 2 bp distance of each other across the 125,748 exome-sequenced individuals from our gnomAD 2.1 dataset, resulting in the discovery of 31,510 MNVs exist within the same codon. When the two variants comprising the MNV were considered together, the resulting functional impact on the protein differed from the independent impacts of the individual variants in nearly 60% of cases (18,714 MNVs; Fig 2a and Supplementary File 1). Among the differing annotations of functional consequence, 405 were “gained” nonsense (neither individual SNV was a nonsense mutation, but the resulting MNV is), and 1,818 were “rescued” nonsense (at least one of the two individual SNVs would create a nonsense mutation, but the resulting MNV does not). Such categories of MNVs have a major impact on variant interpretation, and thus are critical for accurate variant annotation. There was an average of 55.2 variants with altered functional interpretation (including 0.062 gained and 4.42 rescued nonsense) due to MNVs per individual.

**Figure 2.**
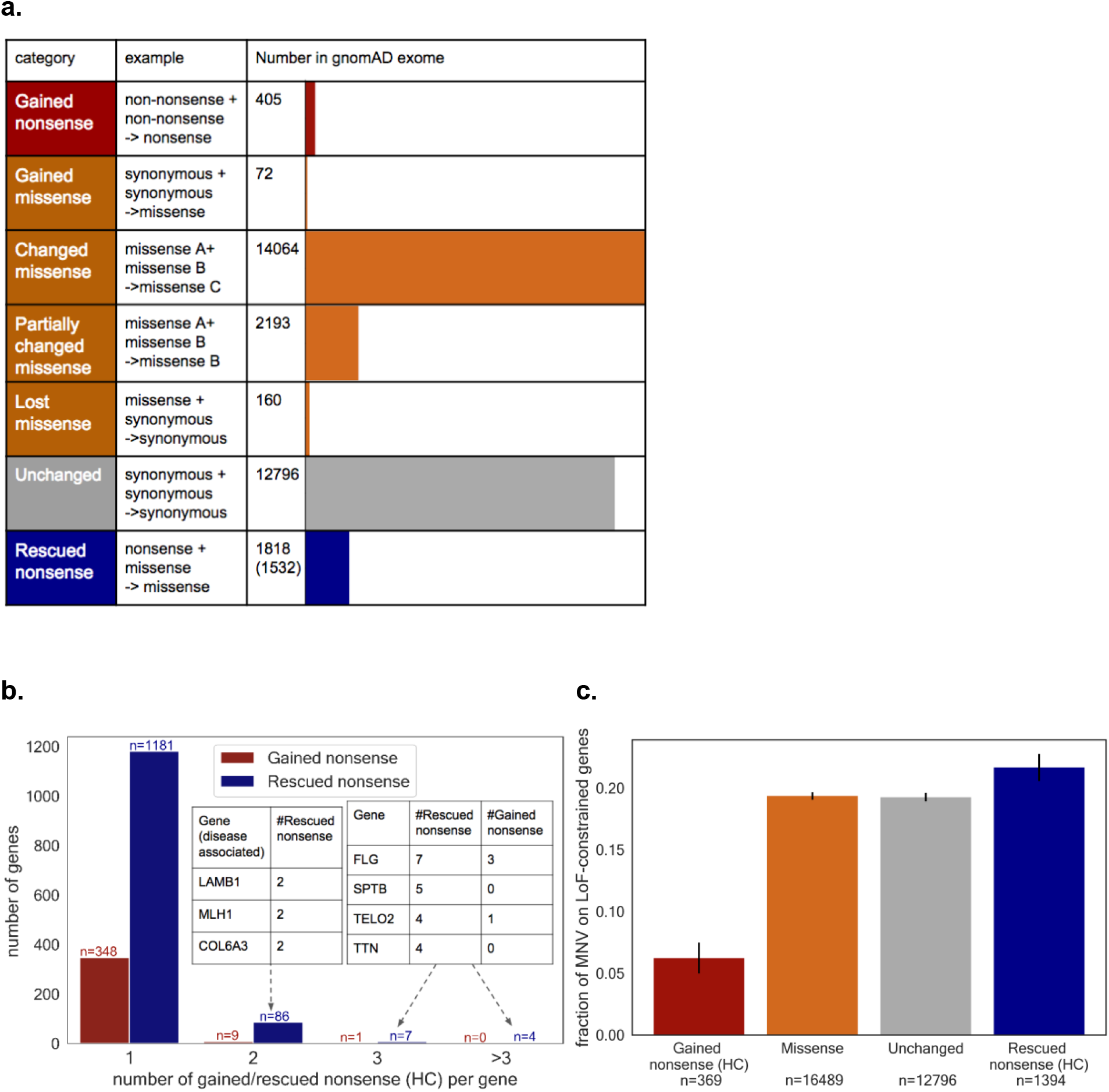
Functional impact of MNVs. **a**, The number of MNVs in the gnomAD exome dataset, per MNV category. Of the 1,818 rescued nonsense mutations, 1,532 are rescued in all individuals that harbor the original nonsense mutation and are used for the analysis in **(b)** and **(c)**. Gained and rescued nonsense MNVs were further filtered to high-confidence (HC) pLoF in **(b)** and **(c). b**, The number of gained/rescued nonsense mutations per gene, and examples of disease-associated genes with 2 or more gained/rescued nonsense mutations. **c**, The fraction of each category of MNV found in a set of 3,941 constrained genes (top two deciles of constraint^22^).

To understand the overall impact of correctly annotating the functional consequence of MNVs in a population-level dataset, we counted the number of gained/rescued nonsense mutations per gene in gnomAD (Fig. 2b, and Supplementary File 1). For rescued nonsense mutations we found 1,532 sites that are rescued in all the individuals with the component variants. A total of 1,594 genes carried gained or rescued nonsense mutations within our dataset, including 41 genes that are disease-relevant (reported by OMIM^23^ or annotated as haploinsufficient by Clingen^24,25^). In addition, the proportion of rescued nonsense mutations of falling in predicted loss-of-function (pLoF) constrained genes (genes with a significant depletion of pLoFs compared to an expectation based on a mutational model^1,26^, defined as LOEUF^22^ decile < 20%) was higher (proportion=0.217) when compared to all the other classes of MNVs (proportion=0.191; Fisher’s exact test, p = 0.0390; Fig 2c and Supplementary Fig. 2). Conversely, gained nonsense mutations are depleted among constrained genes (proportion=0.0623) compared to all other classes of MNVs (Fisher’s exact test, p = 1.40 × 10^−11^). These results suggest a significant enrichment of LoF annotation errors in the absence of MNV annotation.

In order to understand the impact of these variants in clinical applications, we also annotated MNVs in 6,072 sequenced individuals from rare disease families, including 4,275 case samples. This resulted in 16 gained nonsense mutations and 110 changed missense MNVs with high CADD^27^ scores and low frequencies in gnomAD (CADD>20 and <10 individuals in gnomAD; Supplementary File 2). However, after close manual curation, none of the corresponding MNVs were definitively causal variants for the diseases affecting the family, suggesting that MNVs contribute to only a small fraction of total rare disease diagnoses, in line with expectations based on their relative rarity and previous results^2^.

### Genome-wide mutational mechanisms of MNVs

We next turned our attention to understanding the mutational mechanisms underlying the origins of MNVs genome-wide, focusing on whole genome sequence data from 15,708 individuals in the gnomAD v2.1 callset. We considered pairs of high-quality variants in autosomes separated by up to 10 bp, resulting in the assembly of a catalogue of 6,261,326 MNVs including 1,996,125 MNVs within 2 bp distance - an order-of-magnitude increase in size over previous collections.

We considered three established major categories of mutational origins of MNVs with constituent SNVs falling next to each other (“adjacent” MNVs. Fig. 3a), each of which is biased towards certain MNV patterns: (1) combinations of distinct single nucleotide mutation events; (2) replication errors by error-prone polymerase zeta; and (3) polymerase slippage events at repeat junctions. MNVs in the first category are a product of two or more SNVs, which typically occur in different generations and may thus have different allele frequencies. We expect to see an enrichment of CpG transition compared to non-CpG transversion for this class, due to the underlying difference of SNV mutation rate^28–30^. The second category, replication error introduced by DNA polymerase zeta (pol-zeta), is a well known class of replication error that introduces MNVs. Previous studies^10,11–13,31^ have shown that pol-zeta is prone to specific types of replication error, mainly TC->AA, GC->AA, and their reverse complements, with experimental evidence that these MNV patterns occur in a single generation; thus, the constituent SNVs will typically have the same allele frequencies. The third category, replication slippage, is another known mode of DNA replication error^32–34^. This process is especially frequent at sites with repetitive sequence context; previous studies^35–37^ have shown that the insertion and deletion (indel) rate can be up to 10^6^ times higher than the SNV mutation rate at these sites. As shown in Fig. 3a, the combination of an insertion and then a deletion of 2 base pairs can result in an MNV.

**Figure 3.**
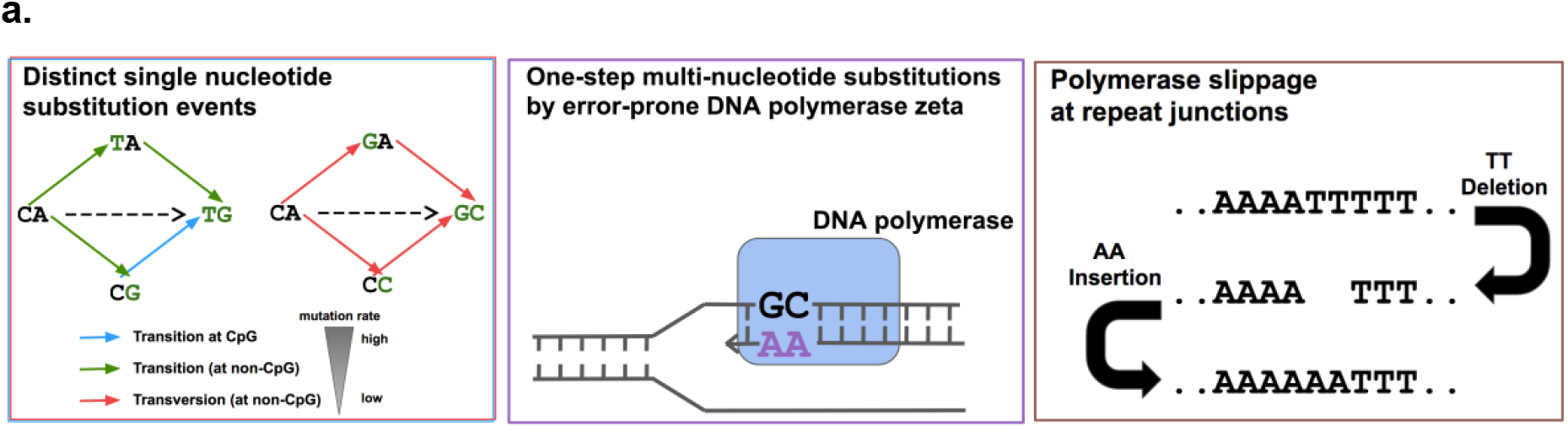
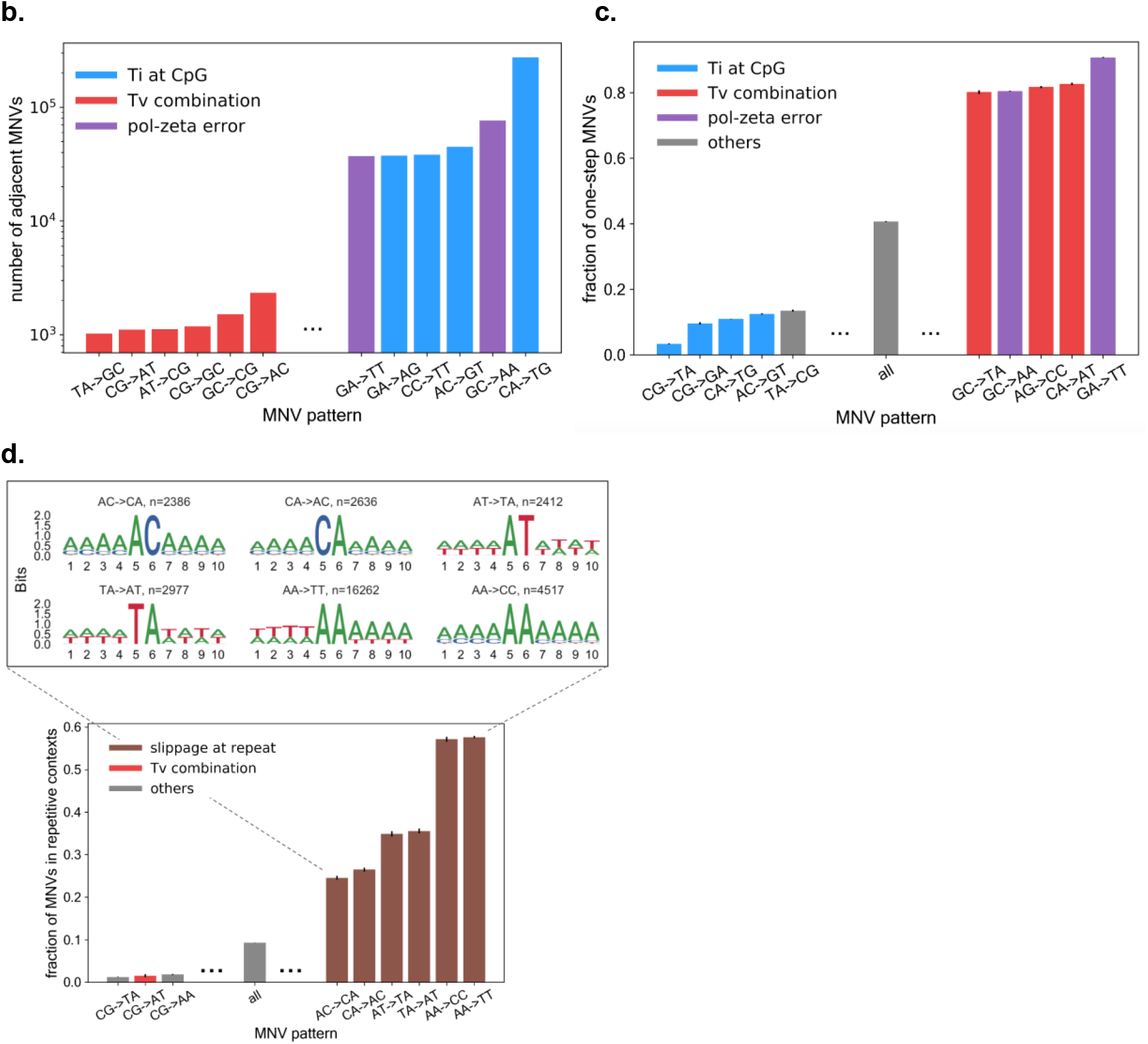
Mutational origins of MNVs. **a**, Three major categories of the mutational origin of MNVs. (Left) A combination of single nucleotide mutational events. Since the baseline global mutation rate is highly different between transversions and CpG and non-CpG transitions, even a simple combination of single nucleotide mutational events could result in a highly skewed distribution of MNVs. (Center) One step mutation caused by error prone DNA polymerases. For this class of MNVs, since the two mutations occur at once during DNA replication, the allele frequency of the two constituent SNVs of the MNV is more likely to be equal. (Right) Polymerase slippage at repeat junctions. Mutation rates are highly elevated in repeat regions, and are therefore likely to cause various complex patterns of mutations, occasionally resulting in MNVs. **b**, The log-scaled number of MNVs per substitution pattern. **c**, The fraction of one-step MNVs per substitution pattern. Error bars represent standard error of the mean. **d**, The fraction of MNVs that are in repetitive contexts, and bits representation^38^ of sequence contexts. MM->NN type MNVs, especially AA->TT, are more likely to fall in repetitive sequence contexts than other MNV signatures. Error bars represent standard error of the mean. Colors in the bars in **(b)** to **(d)** represents the predicted major mechanism of MNVs for each substitution pattern.

We observed the signature of each of these MNV mechanisms in our dataset. First, we calculated the number of MNVs for each MNV pattern (Fig. 3b) and observed that the most frequent MNV pattern is CA->TG substitutions, which are likely to occur as a combination of an A->G transition, followed by a high mutation rate C->T CpG transition. On the other hand, the least frequent MNV pattern is TA->GC substitutions, which occur as a combination of two non-CpG transversions. The 268.5-fold difference (275,240 versus 1,025) of the frequency of MNVs between these two patterns is comparable to the theoretical ratio calculated based on the mutation rate of the component SNVs (471.0-fold), and the overall correlation between the theoretical and observed frequency of each MNV pattern was strong (Pearson correlation r=0.82 with p=2.4 × 10^−4^ in log space; Supplementary Fig. 3).

To investigate the extent of pol-zeta signature, we calculated the number of MNVs in which the gnomAD allele counts of the constitutive single nucleotide variants are equal (following previous methodology^2^), and observed that these “one-step” MNVs are significantly enriched in MNV patterns matching the pol-zeta signature (90.8% for GA->TT, and 80.5% for GC->AA, compared to 40.7% overall; Fisher’s exact test p < 10^−100^; Fig. 3c).

Finally, in order to capture polymerase slippage events, we calculated the fraction of MNVs in repetitive contexts per MNV pattern (Fig. 3d). For the MNV patterns AA->TT and AA->CC, more than 50% of all the MNVs observed were in repetitive contexts. The fractions of the MNV patterns AT->TA and TA->AT in repetitive contexts were also high, exceeding 30% (Fisher’s exact test p < 10^−100^ compared to the 9.25% across all patterns). For all MNV patterns in repeat contexts, we see a significant excess of MNVs compared to the expected number based on a model that assumes MNVs are simple combination of two SNV events (Supplementary Fig. 3). These observations support the role of replication slippage as one of the major drivers of MNVs. Additionally, we did not see a correlation between the frequency of one-step MNVs and the frequency of MNVs in repetitive contexts (the latter was 1.79 fold higher for AA->TT and AA->CC, 0.53 fold lower for AT->TA and TA->AT) for the MNV patterns in the third category, suggesting that multiple slippage events leading to MNV generation can take place either as a single event (i.e. in single generation) or multiple events (i.e. in different generation), or even recurrently.

### Estimation of global mutation rate of MNVs

In order to compare the frequency of three different mechanisms, we quantified the contribution of two single nucleotide variation events vs other replication error modes, such as pol-zeta errors or replication slippage, using a simple probabilistic model. Specifically, focusing on adjacent MNVs, we assigned the MNV frequency for each MNV pattern to be the sum of the probability of two SNV events *(P)* and the probability of other replication error factors (Q), and estimated the *Q* term. In other words, we estimated the divergence of the observed number of MNV sites from the number expected by a simple SNV mutation model (see methods). The resulting estimated proportion of two SNV events and other replication error events is described in Fig. 4a.

**Figure 4.**
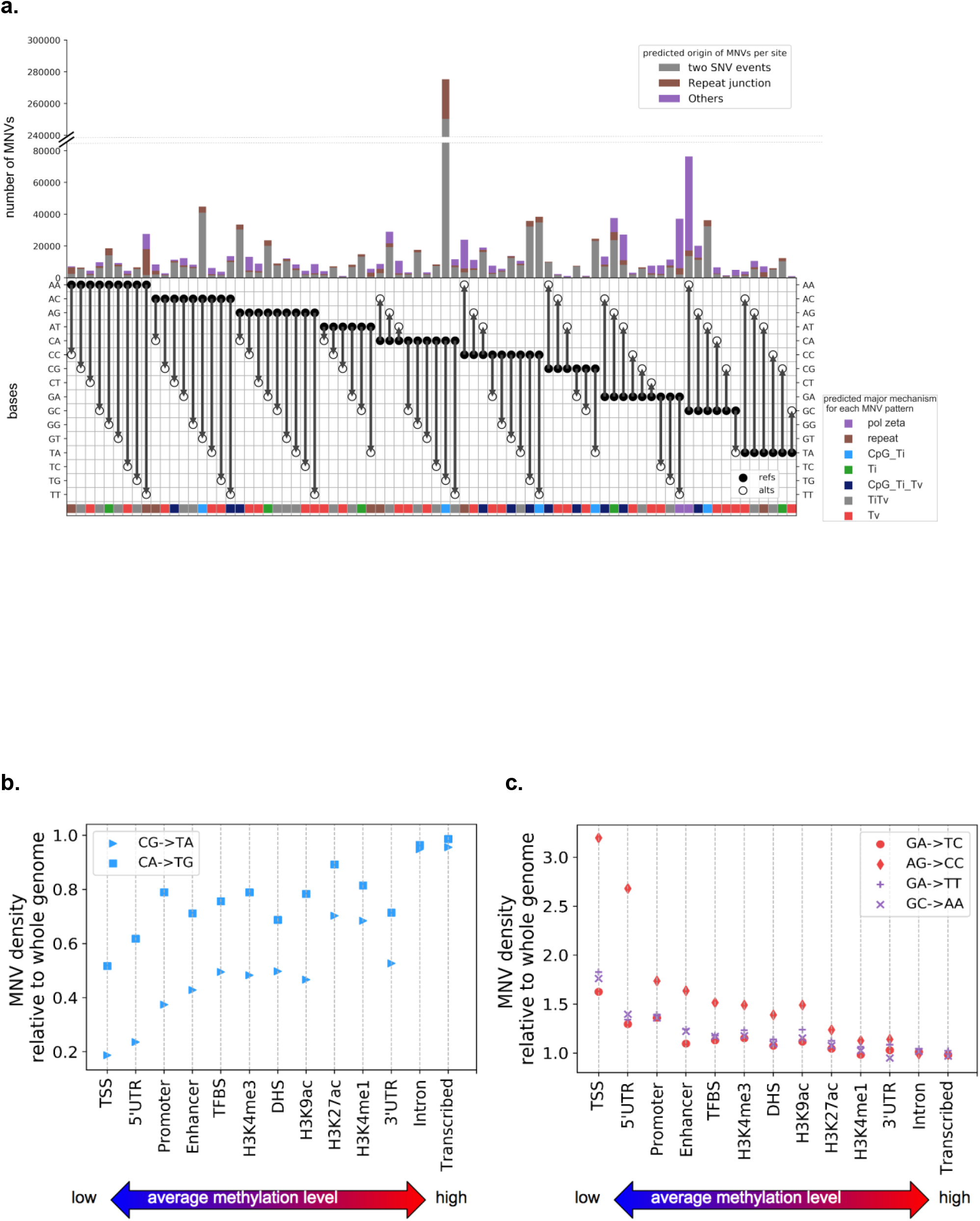
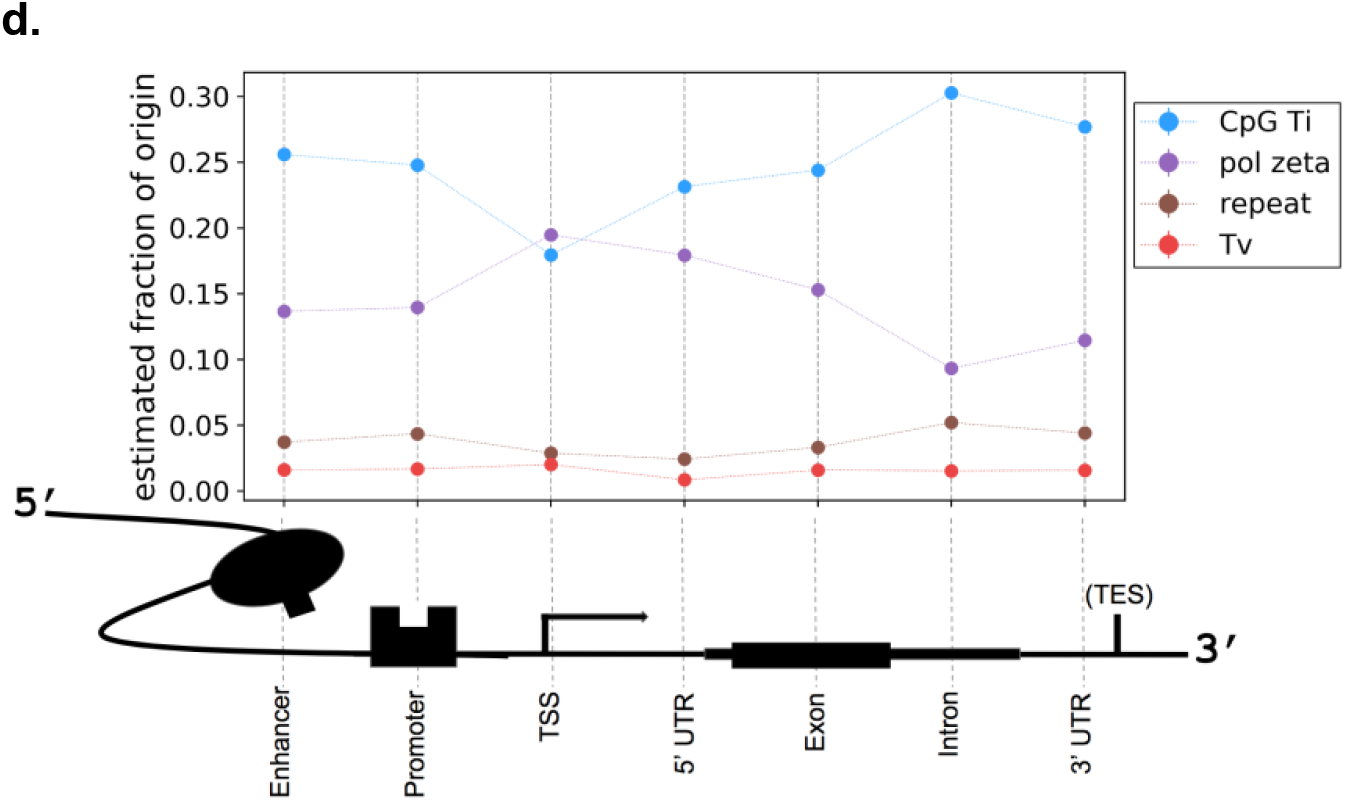
Distribution of MNVs across genome. **a**, The number and the fraction of MNVs per origin, per substitution pattern. Grey are the estimated fraction of MNV originating from two single nucleotide substitution events, brown for polymerase slippage at repeat contexts and purple are the others (presumably mainly replication error by pol-zeta). The colors along the bottom represent the estimated biological origins that dominate MNVs of that specific substitution pattern. **b,c**, MNV density, defined as the number of MNVs per functional annotation divided by the base pair length in the annotation (relative to the whole genome region), ordered by the methylation level of the functional category. **d**, Estimated fraction of MNVs by different origins, per functional category around the coding region.

As expected, the proportion differs substantially from one MNV pattern to another. For example, while 90.1% of CA->TG MNVs appear to be caused by combinations of simple SNV events, the corresponding proportion is 5.42% for GA->TT, 17.8% for GC->AA and 6.36% for AA->TT MNVs. We presume that the lower proportion of two simple SNV events is mainly due to pol-zeta errors for GA->TT and GC->AA, and polymerase slippage for the AA->TT. Since 85% of the overall MNVs were classified as either SNV combination, repeat context, or pol-zeta error at GA->TT or GC->AA, our analysis suggests that these three major categories explain a substantial fraction of MNV events genome-wide, although some possible additional mechanisms with smaller frequencies might exist. These calculations also allow us to estimate the genome-wide mutation rate of MNVs caused by pol-zeta: 1.32 × 10^−10^per 2 bp per generation for GA->TT, and 3.35 × 10^−10^ for GC->AA. Given that there are roughly 1.68 × 10^9^GA pairs and 1.20 × 10^9^GC pairs in the reference human genome, we estimate there are on average 0.22 GA->TT and 0.40 GC->AA mutations per generation (Supplementary File 3).

We also explored the potential mutational mechanisms for MNVs with a greater distance between the component variants (Supplementary Fig. 4,5 and 6), and observed signatures of non-independence of mutation events extending over distances up to 10 bp, with an enrichment of motifs consistent with pol-zeta and polymerase slippage mechanisms for adjacent MNVs (minimum 1.34, maximum 4.10 fold enrichment of one-step MNV, Fisher’s exact test p-value <0.05; Supplementary Fig. 7 and 8). This confirms the presence of mutational mechanisms capable of creating simultaneous mutations separated by considerable distances^16,29,47–49^, although further work will be required to fully characterize the underlying processes.

Overall, our analysis of MNVs in 15,708 whole genome-sequenced individuals supports the previously suggested three major mechanism of MNVs and quantifies the different contribution of each mechanism for different MNV patterns at genome-wide scale.

### MNV distribution across different genomic regions

We next examined how MNV pattern distributions differ between functional annotation categories. We used 13 different functional annotations such as coding sequence, enhancer, and promoter from Finucane *et al*^39^, and the DNA methylation annotation from the Encyclopedia of DNA Elements (ENCODE)^40^, to calculate the number of MNVs that fall into each category (Supplementary Table. 2). MNV density, defined as the number of MNVs observed in each functional category divided by the total length of the genomic interval belonging to each category, is shown in Fig. 4b and c. We found that MNV density of the substitution patterns typically involving CpG transitions is positively correlated with the methylation level (linear regression Pearson correlation r=0.94 for CG->TA and r=0.87 for CA->TG). Conversely, MNV density for non-CpG transversion-related substitution patterns, and the substitution patterns related to pol-zeta slippage, negatively correlates with methylation status (linear regression Pearson correlation r=−0.88 for GA->TC, r=−0.91 for AG->CC, r= −0.89 for GA->TT, and r=−0.92 for GC->AA; Fig. 4b and c).

Finally, we explored the effect of genic context on MNV origins and discovery: we selected the seven major regional annotations around gene coding sequences^41,42^, and calculated the fraction of MNVs likely explained by different mutational origins in each of these regions (Fig. 4d). Across all regions, we found that the MNV signal is primarily dominated by CpG transitions. The fraction of non-CpG transversions and polymerase slippage at repeats were consistently lower than (or nearly equal to) 5% of the overall signal. Pol-zeta signature was not as dominant as CpG transitions, except for at the transcription start site region, which has by far the lowest methylation rate in those seven annotations and is thus expected to have a lower rate of CpG deamination mutations (which are dependent on the methylation of the original cytosine).

Overall, our results suggest that MNV density is highly dependent on the CpG methylation status of the surrounding sequence, and that MNVs that originate from non-CpG transversions or polymerase slippage at repeat junctions are relatively uncommon compared to those driven by CpG transitions or pol-zeta errors. Finally, MNVs that originate from pol-zeta error are the most common class of MNVs in the region close to the transcription start sites of genes, as low methylation levels in these regions result in low levels of CpG transitions.

## Discussion

We analysed 125,748 human exomes and 15,708 genomes and identified 1,996,125 MNVs across genome with constituent variants falling within 2 bp distance, including 31,510 that exist within a codon. We have shown that MNVs represent an important class of genetic variation, and that they have a significant impact on the functional interpretation of genomic data, both at the population and individual level. Although we did not encounter an individual in which an MNV is the likely cause of a rare disease after sequencing 6,072 individuals from rare disease families, we expect that applying our pipeline to larger numbers of disease samples will identify previously missed diagnoses, as has been observed in another study of developmental delay cases^2^.

The large number and high quality of variant calls in the gnomAD database provided increased power for statistical analysis of the three major mutational mechanisms (combinations of independent SNVs; replication errors by pol-zeta; and polymerase slippage at repeat junctions) responsible for the generation of MNVs, and importantly allowed us to estimate the relative contribution of each of these processes.

Our estimates of substitution pattern-specific MNV mutation rate and fraction come with important caveats. Our approach assumes that the local SNV mutation rate is invariant across instances of a specific 3 bp context; however, prior work has shown considerable regional variation in mutation rate across the genome, as well as variation driven by ancestry, environment, and other factors^43–46^. Another important limitation is the lack of confident estimates of insertion and deletion rate as a function of repeat length, which limits the confidence of our estimate of the fraction of polymerase slippage. Future large genome-scale data sets with more accurate insertion and deletion calls, likely involving long-read sequencing data, will be required to improve modeling of insertion and deletion mutations.

One clear feature of our data set was the signature of non-independence of mutational events separated by up to 10 bp, as suggested in various *de novo* studies^16,29,47–49^; further investigation of these clustered mutations, and contextualizing them with known sources of genomic instability, such as homologous recombination^50^ or transposable elements^51,52^, will be informative in exploring the mechanisms of clustered mutations.

The complete list of MNVs identified in gnomAD is publicly available (https://gnomad.broadinstitute.org/downloads), with the allele count annotated for both genome and exome. For the coding regions, we have also annotated the functional consequence of constituent SNVs and MNVs separately, and made the result viewable in an intuitive browser (https://gnomad.broadinstitute.org). Although some fraction of MNVs is missing from this list due to incomplete phasing sensitivity and read coverage, the database provides the most comprehensive set of estimates of MNV allele frequencies to date, valuable for further analysis of mutational mechanisms as well as the interpretation of MNVs in rare disease and cancer genomics^53,54^.

Finally, despite the large sample size of our MNV dataset, the fraction of MNVs that we have observed out of all the possible MNV configurations is still very far from saturating the space of possible MNVs, with only ~0.005% of all possible adjacent MNVs observed in our data (Supplementary Fig. 9, 10). Increasing the number of sequenced individuals in both disease and non-disease cohorts will permit the discovery and determination of the phenotypic impact of an increasingly comprehensive catalogue of variation. This study confirms the importance of incorporating haplotypic phase into these efforts to permit the discovery and accurate interpretation of the full range of human variation.

## Methods

### MNV calling

125,748 human exomes and 15,708 genomes from gnomAD 2.1 callset were used for the analyses (Supplementary Table 3). We used hail (https://github.com/hail-is/hail), an open source, cloud-based scalable analysis tool for large genomic data. For MNV discovery, we exhaustively looked for variants that appear in the same individual, in cis, and within 2 bp distance for the exome dataset and 10 bp distance for the genome dataset, using the hail *window_by_locus* function (i.e. we computationally checked every pair of genotypes within a certain window size, for every individual, to see whether the individual carries a pair(s) of mutation in the same haplotype. See supplementary text for further detail). For trio-based analyses, we expanded the range to 100 bp to obtain a more macroscopic view. Although we performed MNV calling in sex chromosomes for the coding region, we restricted our analysis to autosomes, in order to control for differences in zygosity.

MNV calling in rare disease samples was performed in a similar fashion as in the gnomAD exome dataset. 6,072 rare disease whole exome sequences were curated at the Broad Center for Mendelian Genomics (CMG)^56^ and went through the MNV calling pipeline with the window size of 2 bp distance. The phenotypes observed in the cohort include: muscle disease such as Limb Girdle Muscular Dystrophy (LGMD; roughly one-third of the total), neurodevelopmental disorders, or severe phenotypes in eye, kidney, cardiac or other orphan diseases (Supplementary File 2).

### MNV filtering

In the gnomAD MNV analysis, MNVs for which one or both of their components have low quality reads were filtered out. Specifically, we only selected the variant sites that pass the Random Forest filtering, resulting in acceptance of 53.3% of the initial MNV candidates (Supplementary Fig. 11). For the exome dataset, we also applied adjusted threshold criteria (GQ ≥ 20, DP ≥ 10, and allele balance > 0.2 for heterozygote genotypes) for filtering individual variants. For each MNV site, we annotated the number of alleles that appear as MNV, as well as the number of individuals carrying the MNV as a homozygous variant. The distribution of MNV sites that contain homozygous MNVs is shown in Supplementary Fig. 12. We also collapsed the MNV patterns that are reverse complements of each other, after observing that the number of MNVs are roughly symmetric (Before collapsing, the ratio of each MNV pattern to its corresponding reverse complement pattern was mostly close to 1, with 0.91 being the lowest and 1.05 being the highest for adjacent MNVs) (Supplementary Fig. 13). All the MNV patterns in the main text and figures are equivalent to their reverse complement, and we do not distinguish them.

For the rare disease cohort, since our motivation was to find a definite example where an MNV is acting as a causal variant for a rare disease with severe phenotype rather than obtaining the population level statistics, we did not apply site and sample-specific filtering, as opposed to the gnomAD MNV analysis. Instead of being computationally filtered by read quality, the 129 putative MNVs (16 gained nonsense mutations, 110 changed missense with high CADD score and low gnomAD MNV frequency, and 3 gained missense) went through manual inspection by the analysts at the Center for Mendelian Genomics (CMG) at the Broad Institute^56^, after annotating the affected gene. Specifically, all the variants were checked manually under the criteria below:

- Whether the gene affected is constrained in the gnomAD population
- Whether the case has already been solved with other causal variant
- Whether the MNV looks real in the Interactive Genome Browser (IGV)^57^
- Whether the MNV is in the proband and, if applicable, the segregation pattern of the MNV
- Whether the known function of the gene affected matches the patient phenotype

MNVs were filtered out if they failed one or more of the criteria above. These results suggest that MNVs explain only a small fraction of undiagnosed genetic disease cases, consistent with their overall frequency as a class of variation, and with prior work in large disease-affected cohorts^2^. The summary for MNV analysis in rare disease cohort is also available at Supplementary File 2.

### Analysis of phasing sensitivity

In order to compare the phasing information derived from different methods (read-based and trio-based), we took an approach of comparing the relative phase (binary classification of whether two SNVs of MNV are in the same haplotype or not), as shown in Supplementary Table 4. We investigated the heterozygous variant pairs whose phasing information is not provided by the trio-based phasing and observed that majority (83.5%) of the cases reflected both parents carrying a heterozygous variant, a scenario where trio-based phasing is inherently uninformative. We also investigated the heterozygous variant pairs whose phasing information is not provided by the read-based phasing. Specifically, unphased pairs tend to have either low or high read depth (odds ratio =3.20, Fisher’s exact test p < 10^−100^ for low, and odds ratio =2.33, Fisher’s exact test p < 10^−100^for high read depth; Supplementary Table 1), consistent with our previous understanding that an excess of reads can lead to involvement of erroneous reads and thus reduce the confidence of phasing of HaplotypeCaller^58^ (as well as the lack of the number of reads reduces the calling rate).

### Analysis of functional impact in coding region

We focused on the coding region of the canonical transcript of genes and examined the codon change and their consequence for all the MNVs that fall in a single codon. When comparing with population level constraint, for each MNV, we annotated the constraint metric (LOEUF^22^) of the gene whose protein product is affected. For rescued nonsense mutations, we took only the ones are rescued in all the individuals with the component variants (i.e. we excluded the ones whose allele count of MNVs are not equal to the allele count of the SNV that introduces a nonsense mutation), resulting in 1,532 out of 1,818 rescued nonsense mutations. We next used Loss-Of-Function Transcript Effect Estimator (LOFTEE^22^) in order to exclude the nonsense mutations that are not likely to affect the protein function. This resulted in 369 high-confidence (HC) gained nonsense mutations and 1,394 HC rescued nonsense mutations, which were used for the population-level constraint analysis. In addition, we stratified the gene sets by core essential/non-essential genes from CRISPR/Cas knockout experiments^59,60^ as an orthogonal indicator of gene constraint (Supplementary Fig. 2.).

Although theoretically a combination of insertions and deletions of different lengths could also change the individual consequence of the variants (for example, an insertion of length 4 followed by a nearby deletion of length 1 results in a insertion of length 3, which restores the codon reading frame), we focused on SNV combinations and did not try to identify such class of variants in this work. Also, we did not include and correct for MNVs consisting of three SNVs in a single codon in the analysis of functional impact in coding region, since the number and frequency of such MNVs are significantly low (231 in total, with 5 newly gained nonsense, but no re-rescued or re-gained nonsense. 0.220 in total per person). The full list of such MNVs are available as a separate file at (https://gnomad.broadinstitute.org/downloads).

### Defining one-step MNVs and MNVs in repetitive contexts

A one-step MNV was defined as a MNV for which the allele count of both SNVs that make up the MNV is the same and close to the allele count of the MNV itself. We also compared the allele count of constituent SNVs (AC1 and AC2) with the allele count of the corresponding MNV (AC_mnv), and observed that the majority of one-step MNVs we discovered have AC_mnv/AC1 >0.9 (Supplementary Fig 14). Therefore we expect the false discovery rate of one-step MNVs (misclassifying the MNV whose AC1 and AC2 are equal just by chance) to be limited. The full distribution of all the allele counts are shown in Supplementary Fig 15.

Repetitive sequences are defined by taking the +/− 4 bp context of the MNV and setting the threshold manually, by looking at the distribution of repeat contexts around all the MNVs (Supplementary Fig. 16, 17). Specifically, for adjacent MNVs, a sequence is defined as repetitive if

- there is a >6 bp mononucleotide repeat, for both reference and alternative +/− 4 bp context, or
- there is a ≥6 bp dinucleotide repeat, for both reference and alternative +/−4 bp context

This threshold was set so that the number of MNVs with equal or higher repeats would be less than 5% of the total. Also, the estimated mutation rate under those repetitive contexts is thought to be orders of magnitude higher than the background MNV mutation rate originating from the combination of two SNV events.

### Calculating the proportion of MNVs per biological origin

We calculated the proportion of MNV per biological origin by comparing the observed number of MNVs (that are not in repetitive contexts) with the expected number of MNV under single nucleotide mutational model.

Specifically if we simply hypothesize most of the MNV are combination of two single nucleotide substitution events, we can estimate the relative probability of MNV event per substitution pattern. For example, probability of observing a CA to TG MNV in a single individual, single site (*p*(*CA* → *TG*)) is proportional to *p*(*CA* → *TA*) · *p*(*TA* → *TG*) + *p*(*CA* → *CG*) · *p*(*CG* → *TG*), and probability of TA to GC MNV (*p*(*TA* → *GC*)) is proportional to *p*(*TA* → *GA*) · *p*(*GA* → *GC*) + *p*(*TA* → *TC*) · *p*(*TC* → *GC*). Former equation involves the product of transition at CpG, while both term of the latter are product of transversion at non-CpG, which works as a reasonable explanation of the frequency difference of those two MNV patterns.

Using the same principle (and accounting for reference base pair frequency, population number and global SNV mutation rate defined by 3 bp context^26^, we first constructed a “null model” of MNV distribution. In reality, this null model does not represent the real distribution we observe, due to biological mechanisms that introduce MNV. Therefore, we allowed additional factor q, that denotes the mutational event where two SNVs are introduced at the same time. For the example of *p*(*CA* → *TG*), we model this probability to be proportional to *p*(*TA* → *GA*) · *p*(*GA* → *GC*) + *p*(*TA* → *TC*) · *p*(*TC* → *GC*) + *q*(*CA* → *TG*), and try to estimate the *q* term, which corresponds to the proportion of MNVs that are explained by non-SNV (and non-repeat) factor. Further details are explained in the supplementary text (section “**Models and assumptions for calculating the proportion of MNV per biological mechanism**”).

In addition, for each of MNV pattern, we annotated the predicted major mechanism for each MNV pattern in the following order:

1. “pol-zeta”, for the patterns known as polymerase signature (GA->TT and GC->AA)
2. “repeat”, for the patterns whose fraction of MNVs in repeat contexts are higher than 20%
3. One of “CpG_Ti”, “Ti”, “CpG_Ti_Tv”, “TiTv”, “Tv”, based on possible combinations of single nucleotide mutational processes. For example, “CpG_Ti” is when transition in CpG combined with another transition can occur in the mutational processes (Supplementary File 3)

### Estimation of the global MNV rate per substitution pattern

In order to estimate the global MNV mutation rate for adjacent MNVs, as well as the mutation rate per MNV pattern, we first focused the number of one-step MNVs, assuming that there are no recurrent mutations and therefore the allele frequency of constituent SNVs are equal if and only if if originates from an MNV event in a single generation. In this section, we will simply write one-step MNV of distance 1 bp (i.e. adjacent) as MNV.

We then calculated the global MNV mutation rate under the Watterson estimator model, as in Kaplanis et al^2^. Specifically, we divided the number of MNV sites by the number of SNV sites in our gnomAD dataset, and scaled by the global single nucleotide mutation rate identified in previous research (1.2 × 10^−8^), which yielded 2.94 × 10^−11^per 2 bp per generation. This is roughly two thirds of the estimation provided by the Kaplanis et al^2^ using trio data, slightly smaller presumably due to differing filtering method. Next, In order to get the mutation rate per 2 bp for each of the MNV patterns, we simply scaled the global MNV mutation rate described above by the number of reference 2 bp and the coverage difference. The full data for all the 78 patterns is shown in Supplementary File 3. Further details are explained in the supplementary text (section “**Models and assumptions for estimation of the global MNV rate per substitution pattern**”).

### Functional enrichment

13 functional annotations were collected from Finucane et al^39^ as a bed file (which originates from database such as ENCODE, Roadmap^61^ and UCSC genome browser^62^.) For the methylation data, we collected the genome methylation level from ENCODE, and calculated the fraction of methylated CpG out of all the CpGs in the region, and ordered by the fraction (Supplementary Table 2).

MNV density calculation was performed under the null hypothesis that, the number of MNV of type *WX → YZ* we observe in an arbitrary genomic interval is proportional to the number of *WX* in the interval. Specifically, the MNV density of *WX → YZ* in interval *I* is defined as *D*(*WX → YZ|I*) = *N*(*WX → YZ|I*) / *N*(*WX*|1), where *N*(*WX* → *YZ | I*) *is* the number of MNVs of *WX → YZ*, and *N*(*WX|I*) *is* the number of *WX* in the reference genome we observe in that specific genomic interval. We then normalized the density by dividing by *D*(*WX → YZ| I* = *whole genome*) *for* scaling purpose (i.e. *D*(*WX → YZ|I*) = *k* means that the probability of observing a mutation of *WX → YZ* given a sequence context of *WX* is *k* times higher in genomic functional category *I* than the overall genome.)

For estimating the fraction of MNVs per origin, we took a thresholding approach and defined four MNVs (CA->TG, AC->GT, CC->TT, and GA->AG) as CpG signal, two (GC->AA, GA->TT) as pol-zeta, five as repeat (AA->CC, AA->TT, TA->AT, AT->TA, CA->AC, AC->CA) and transversion (TA->CG, AC->GT, CA->TG, CG->GA, CG->TA) signal (and left all the other 78-(4+2+5+5)=62 patterns as “others”, in order to highlight the strongest signals) based on the result from Fig. 3. The fraction of MNVs per origin is then defined simply as the number of MNVs that fall into that pattern divided by all the MNVs, in the genomic interval. The coverage difference per interval was as small as negligible (Supplementary Table 3).

### Data availability

The list of coding MNVs in gnomAD exome are available at gs://gnomad-public/release/2.1/mnv/gnomad_mnv_coding.tsv (tab separated file). The coding MNVs consisting of three SNVs in a single codon is available as a separate file at gs://gnomad-public/release/2.1/mnv/gnomad_mnv_coding_3bp.tsv.

The list of all the MNVs in gnomAD genomes are available at gs://gnomad-public/release/2.1/mnv/genome/gnomad_mnv_genome_d{i}.tsv.bgz (tab separated file, compressed. Replace {i} (0<i<11) with the distance between two SNVs of MNV.), or gs://gnomad-public/release/2.1/mnv/genome/gnomad_mnv_genome_d{i}.ht (hail table. Replace {i} (0<i<11) with the distance between two SNVs of MNV.).

Explanations for each column in each file can be found at gs://gnomad-public/release/2.1/m nv/mnv_readme.md.

All the files above are also available at the download page of the gnomAD browser (https://gnomad.broadinstitute.org/downloads).

### Code availability

The code used in the study is available at https://github.com/macarthur-lab/gnomad_mnv.

## Acknowledgements

We would like to thank the many individuals whose sequence data are aggregated in gnomAD for their contributions to research, and for making this work possible. The results published here are in part based upon data: 1) generated by The Cancer Genome Atlas managed by the NCI and NHGRI (accession: <u>phs000178.v10.p8</u>). Information about TCGA can be found at http://cancergenome.nih.gov, 2) generated by the Genotype-Tissue Expression Project (GTEx) managed by the NIH Common Fund and NHGRI (accession: <u>phs000424.v7.p2</u>), 3) generated by the Exome Sequencing Project, managed by NHLBI, 4) generated by the Alzheimer’s Disease Sequencing Project (ADSP), managed by the NIA and NHGRI (accession: <u>phs000572.v7.p4</u>). We would like to thank the Hail team for developing tools essential for the large scale computation in this work. We would like to thank the analysis team of the Broad’s Rare Disease Group for their manual inspection of MNVs in rare disease cohorts. This work was funded by NIDDK U54 DK105566, NIGMS R01 GM104371, and NHGRI UM1 HG008900-01. KJK was supported by NIGMS F32 GM115208. QW was supported by the Nakajima Foundation Scholarship. We have complied with all relevant ethical regulations.This study was overseen by the Broad Institute’s Office of Research Subject Protection and the Partners Human Research Committee, and was given a determination of Not Human Subjects Research. Informed consent was obtained from all participants.

