## Supplementary Information for "Landscape of multi-nucleotide variants in 125,748 human exomes and 15,708 genomes"

**Supplementary materials for “Landscape of multi-nucleotide variants in 125,748 human exomes and 15,708 genomes”**

### Supplementary methods

#### (MNV calling workflow)

Using hail, we loaded the gnomAD data as a matrix table, pre-filtered the variants that are homozygote (hom) of reference allele (ref), applied the *window_by_locus* function to call MNVs, and used *aggregate* function to obtain the site level information. Our pipeline performs the equivalent calculation and outputs the same result as the python-like pseudo-code below, with orders of magnitude faster computational time compared to running a python script in local, achieved by cloud based parallel computing.

pseudo code:
let the window size be *k*, and a snp in position *i* be *snp_i*

```
for all position i in the genome:
        for j in [i+1, ... , k]:
                for every sample s carrying non-ref allele in both snp_i and snp_j:
                if (both snp_i and snp_j is hom):
                        hom_snp_cnt_{s,i,j} = 1
                elif (only one of snp_i or snp_j is hom):
                        het_snp_cnt_{s,i,j} = 1
                elif ((both snp_i or snp_j is het)
                      & (snp_i and snp_j are phased in a same haplotype)):
                        het_snp_cnt_{s,i,j} = 1
```

Allele counts for each site can be aggregated as

$$hom\_snp\_cnt_{i,j} = \sum_s hom\_snp\_cnt_{s,i,j}$$

$$het\_snp\_cnt_{i,j} = \sum_s het\_snp\_cnt_{s,i,j} \; .$$

For the genome data analysis, the number of MNV (or MNV count) in the main text denotes the total number of unique *(i, j)* pairs where $hom\_snp\_cnt_{i,j} + hom\_snp\_cnt_{i,j} > 0$ (i.e. the number of unique MNV sites). We took a slightly different approach to inferring MNV count for the exome analysis to focus on the functional consequence, by counting the number of MNV existing fully within a codon of a canonical transcript. We note that the number of unique MNV sites (31,133 sites) for exome coding regions is slightly lower than the MNV count (31,510 counts) we provide in the main text, as some MNVs affecting more than one canonical transcripts.

We set $k = 3$ for gnomAD exome and rare disease analysis, $k = 10$ for non-coding, and $k = 100$ for phase sensitivity analysis. The code above assumes no multiallelic variants for simplicity, but in reality we inspected multiallelic variants as well. This can be simple written by replacing *snp_i* with *snp_i_1*, *snp_i_2*, ... *snp_i_n* when there are n different snps at position *i*, and re-writing the 4th line as `*for all the SNP pairs (snp_i_*,snp_j_*)*` by letting "*" denote *1, 2, ... n.)*. This code assumes diploid status for the analyzed individual, but expanding to haploid status can be easily achieved by omitting the process of inspecting phase. We called MNVs in sex chromosomes for the exome and rare disease analysis. For the genome-wide analysis, we did not call the homozygotes whose phasing information is not assigned. We also saved the metadata such as the result of quality control (QC) for each site and sample, and chose to filter the variants or not depending on the study purpose.

**(Models and assumptions for calculating the proportion of MNV per biological mechanism)**

The fraction of non-SNV contribution for each MNV pattern, and its global probability were calculated as follows (note: in this section A or C is simply used according to the alphabetical order without any specification, and therefore does not mean adenine or cytosine):

1. We first took the number of 4 base pair contexts in the reference genome, as well as the median coverage of the region and defined them as follows:

   $N(ABCD)$:= number of instances of the base pair context $ABCD$in the reference genome

   $c(x)$ := median coverage at position $x$

2. We know that the expected rate of detecting a variant is correlated to coverage at the site. We write this as $f(c(x))$.

For the calculation, we empirically approximated this relationship as $f(c) = tanh(c/25)$

3. Let the base line mutation rate defined by 3 bp context be $p_0(ABC \to ADC)$.Considering the coverage difference, the actual probability that we observe that change at the position x, where the 3 bp context of x is $ABC$ is

$$p_x(ABC \to ADC) = p_0(ABC \to ADC) \cdot f(c(x)).$$

4. For a specific type of MNV $ABCD -> AEFD$ with $B$ at position $x$, we calculate the relative probability of this happening as

$$p_x(ABCD \to AEFD) = p_x(ABC \to AEC) \cdot p_{x+1}(ECD \to EFD) + p_x(BCD \to BFD) \cdot p_{x+1}(ABF \to AEF)$$

If we assume the coverage of the bases next to each other is almost identical$f(c(x)) \approx f(c(x+1))$, then

$$p_x(ABCD \to AEFD) = \{p_0(ABC \to AEC) \cdot p_0(ECD \to EFD) + p_0(BCD \to BFD) \cdot p_0(ABF \to AEF)\} \cdot f^2(c(x))$$

5. The overall fraction of adjacent MNV of $ABCD -> AEFD$ can be calculated by integration over all the positions:

$$p(ABCD \to AEFD) = \sum_y p_y(ABCD \to AEFD),$$

where y are all the position across genome with $B$ followed by $C$ as the reference sequence.

6. The overall fraction of MNV of *$BC$* -> *$EF$*, where * can be arbitral of base pair (A, C, G, or T), can be calculated by averaging over all the base patterns:

$$p(*BC* \to *EF*) := p(BC \to EF) = \sum_{a,d} p(aBCd \to aEFd)$$

7. In practice, if we write the average of $f^2(c)$for 4 bp pattern $ABCD$ as $cat(ABCD)$ (standing for "Coverage Adjustment Term"), the number of 4 bp count of ABCD in the reference genome as $N(aBCd)$), and the relative mutation probability for a single site as

$p_0(ABCD \to AEFD) = p_0(ABC \to AEC) \cdot p_0(ECD \to EFD) + p_0(BCD \to BFD) \cdot p_0(ABF \to AEF)$ then we can write the equation in 6. using summary statistics only, without actually scanning through the genome:

$$p(BC \to EF) = \sum_{a,d}\{p_0(aBCd \to aEFd) \cdot N(aBCd) \cdot cat(aBCd)\},$$

8. Now we have the probability matrix of 16x16 $:= P$, where each row denotes the reference and each column denotes the alternative alleles, and the entry corresponds to the relative probability of observing each type of MNV in the human genome. Let the observed MNV matrix be $M$ (in practice, we let $M$ be the number of MNVs that are not in repeat contexts), then we construct the "null matrix" $N$ by multiplying the probability matrix with constant $k$:

$$N = k \cdot P$$

The approximation to estimate the parameter $k$ will be discussed in 10. This null matrix, by definition, is the expected number of MNV for each MNV pattern, assuming single nucleotide substitution process, whose probability is defined by the 3 bp context, is the only driver of MNV. However, as seen in previous studies and our study, this overlooks some important MNV mechanisms such as polymerase zeta error and polymerase slippage at repeat junctions.

9. In order to include additional slippage / polymerase zeta contribution into the model, we added the "other contribution" term $q_0$, which depends only on the 2 bp context. Then the equation can be re-written as

$$p'_x(ABCD \to AEFD) = \{p_0(ABC \to AEC) \cdot p_0(ECD \to EFD) + p_0(BCD \to BFD) \cdot p_0(ABF \to AEF) + q_0(BC \to EF)\} \cdot f^2(c(x)).$$

Then following the same derivation step,

$$p'(BC \to EF) = \sum_{a,d}\{(p_0(aBCd \to aEFd) + q_0(BC \to EF)) \cdot N(aBCd) * cat(aBCd)\}$$
$$= \sum_{a,d}\{p_0(aBCd \to aEFd) \cdot N(aBCd) * cat(aBCd)\} + q_0(BC \to EF) \cdot N(BC) \cdot cat(BC)$$

10. Now we have the matrix $P'$, and can let the null matrix be exactly equal to the observed matrix by letting $P' \cdot k = M$. However, this equation has $16 \cdot 9 + 1$ variables ($16 \cdot 9$ non zero element of matrix plus $k$), with only $16 \cdot 9$ equations. Therefore, we manually set the parameter to make this calculation possible. Specifically, we hypothesize that "for the most common MNV pattern CA->TG, which we think is primarily driven by increased single base mutation rate by CpG methylation, there should be no significant extra factor contribution other than polymerase slippage at repeat junction, and therefore $q_0$ is approximately zero". Specifically, we set $q_0(CA \to TG)$ to be 0, and calculated k and all the other $q_0$ terms:

$$k = m(CA \to TG) \,/\, p(CA \to TG)$$
$$q_{(BC \to EF)} = q_0(BC \to EF) \cdot N(BC) \cdot cat(BC) = m(CA \to TG) \,/\, p(CA \to TG)$$

The resulting matrix $Q$ where $Q_{i,j} = q_{i \to j}$ is used for the estimation of non spontaneous factor contribution. Specifically, if we write $P' = P + Q$, then the fraction $\frac{Q_{i,j}}{(P_{i,j} + Q_{i,j})}$ directly provides the estimation of relative non-SNP contribution. Note that, for the calculations above, since the SNP mutation rates are typically in the order of $10^{-7}$ or lower, any higher order term was approximated as zero (i.e. we are assuming that there are no recurrent mutations).

11. In practice, mainly because of the fact that the mutation rate is not uniform across genomes even given the 3 bp context, the model produces some of the $q_0(i \rightarrow j)$ to be negative ( i.e. we overestimate the underlying mutation rate by SNP combination) (Supplementary Fig 18). For such cases, since the fraction of overestimation is relatively small (less than 5% for most of the cases), we simply replaced the value with zero. Also, presumably because we set a manual threshold for the repeat context definition although in reality we know that the mutation rate increases as a function of repeat number (or sequence non-complexity in general), we might be underestimating the number of MNVs that originate from polymerase slippage at repeat contexts. Therefore, the fraction of "other" term should be interpreted carefully for the cases such as CC->AA, where both the fraction of repeat and others are high (i.e. the repeat count should rather be interpreted as the lower bound).

We also tried to incorporate additional complexity to the mutational model in two ways. First, we calculated the number of mutation sites in the gnomAD dataset per methylation status bin, and let the mutation rate differ as the function of methylation state (supplementary fig 19, (a) and (b)). Second, we expanded the sequence context model to 7 bp context rather than 3 bp, and calculated the mutation rate as a function of 7 bp context (supplementary fig 19, (c) and (d)). In both cases, the result was quite similar to the baseline case, suggesting that 3 bp context alone explains significant amount of MNV frequency.

In addition, we compared two models to adjust for coverage: the first assuming that the probability that each SNV of the MNV is detected is independent (therefore we adjust by the factor $f^2$), and the second assuming that the correlation between the detection of first SNV and the second is exactly equal to 1 (i.e. If we miss the first SNV by sequencing error due to low coverage, we also miss the second SNV that are next to it. Therefore we adjust by the factor $f$, without squaring). Although these two models differ in their assumptions, the difference was minor (>99% consistency in the final count).

Lastly, although theoretically the rate of MNV creation by SNV combination in a single generation is at most in the order of $10^{-15}$ ($10^{-8} \cdot 10^{-7}$in the case of non-CpG transition followed by a CpG transition), in practice we observe several orders of magnitude higher mutation rate overall. Our model takes relatively strong assumption that there is a "general" constant factor that pushes up the MNV mutation rate for all the MNV patterns. Further experiments to deepen our understanding of human genome mutation rate will be required to improve the assumptions made in our study.

**(Models and assumptions for estimation of the global MNV rate per substitution pattern)**

When assuming a fixed effective population size, no recurrent mutations, and the uniformity of the mutation rate across genome, the single base pair mutation rate $\mu_{SNP}$is known to be approximated as proportional to the number of the sites where mutations were observed (=$N(SNP)$).

$$\mu_{SNP} \propto N(SNP) = c_0 \cdot N(SNP)$$

In order to expand this basic model to MNV rate calculation globally and per MNV patterns, we expanded the model in two ways based on previous work (Kaplanis *et al*[2]) method.

1. Under the assumption that there is no recurrent mutation and therefore the allele frequency of constituent SNPs are equal if and only if originates from an MNV event in a single generation, the global mutation rate of adjacent MNV can be written using $N(MNV)$ that denotes the number of 2 bp sites where MNV were observed:

$$\mu_{MNV} \propto N(MNV) = c_0 \cdot N(MNV)$$

(And the constant factor $c_0$ is same for two equations above, because of the fixed population size assumption and the uniform reference genome)

Since we know the SNP mutation rate from previous researches ($1.2 \cdot 10^{-8}$), we can simply calculate the global MNV mutation rate as

$$\mu_{MNV} = \mu_{SNP} \cdot N(MNV) \,/\, N(SNP),$$

Where $N(SNP)$ is the number of SNP sites.

In our data, this calculation resulted in the global MNV mutation rate of

$$\mu_{MNV} = 1.2 \cdot 10^{-8} \cdot 1257972 \,/\, 205390268 = 2.94 \cdot 10^{-11}.$$

2. Under additional assumption that recurrent mutation is negligible for every 2 bp context (assuming that, although we know the single nucleotide mutation rate of CpG is different from other 2 bp context, the difference is minor compared to the MNV mutation rate.), we can compare the MNV mutation rate for different substitution patterns by letting the factor $c_0$ to be proportional to the number of 2 bp count in the reference genome, rather than being a constant. We also account for slightly different coverage per 2 bp context, and write it as $cat(XY)$ (standing for "Coverage Adjustment Term", as in the previous section, but this time defined by 2 bp rather than 4 bp context). Then, by replacing the $c_0$ with $N(XY) \cdot c$, and letting the mutation rate of XY->ZW given a sequence context XY per generation to be $\mu_{XY \to ZW}$, we can write the mutation rate as:

$$\mu_{XY \to ZW} = N(MNV_{XY \to ZW}) \cdot N(XY) \cdot c \cdot cat(XY)$$

Here, the newly appeared constant term c is assumed to be uniform across different 2 bp context, and also when calculating the global MNV mutation rate. Specifically, if we use the term * to denote arbitral base pair (either A, C, G, or T), the global MNV mutation rate calculated in the 1 can be rewritten as:

$$\mu_{MNV} = 2.94 \cdot 10^{-11} = \mu_{** \to **} = N(MNV_{** \to **}) \cdot N(**) \cdot c \cdot cat(**),$$

Where $N(MNV_{** \to **})$ is identical to $N(MNV\ )$ in 1., $N(**)$, the number of 2 bp count in the reference genome, is equal to the number of reference genome length minus one, and the $cat(**)$ can be easily calculated from the median coverage of the entire genome.

With this, for all the 78 patterns of reference and alternative 2 bp, we can calculate the mutation rate as a function of $N(MNV_{XY \to ZW}) \cdot N(XY)$ and $cat(XY)$ (i.e. the number of MNV site of pattern XY->ZW, the number of 2 bp count of XY in the reference genome, and the mean of media coverage of XY):

$$\mu_{XY \to ZW} = N(MNV_{XY \to ZW}) \cdot N(XY) \cdot c \cdot cat(XY)$$
$$= N(MNV_{XY \to ZW}) \cdot N(XY) \cdot \{\, \mu_{** \to **} \,/\, (N(MNV_{** \to **}) \cdot N(**) \cdot cat(**))\} \cdot cat(XY)$$

Finally, comparison with the *de novo* MNV rate from trio exome sequencing data (Kaplanis et al) was performed as follows. Kaplanis et al estimated the global MNV mutation rate for the MNV of distance 1 to 20 bp to be $1.78 \cdot 10^{-10}$per base pair per generation. In order to scale this to our unit, which is MNV mutation rate restricting to adjacent MNVs (MNV of distance 1 bp), per 2 bp per generation, we divided the number by 2, and further multiplied by the fraction of adjacent MNVs out of all the MNVs of distance 1 to 20 bp (both restricting to one-step MNV) they have discovered. Specifically, this resulted in $1.78 \cdot 10^{-10} / 2 \cdot \{6606 \,/\, (6606 + 1568 + 4802)\} = 4.53 \cdot 10^{-11}$, where 6606, 1568 and 4802 are the number of MNVs of distance 1 bp, 2 bp, 3-20 bp they have discovered. This is $(4.53 \cdot 10^{-11}) \,/\, (2.94 \cdot 10^{-11}) = 1.54$ times higher than the estimation provided by our analysis. We assume one of the main reasons for this discrepancy is the fact that the model does not take recurrent mutations into account, and would thus miss some of the MNV events followed by a single base pair mutation event, which might be particularly prevalent when the MNV results in CpG creation. Also, different MNV calling methods, as well as filtering criteria, are likely to contribute to the difference. Since we have no phase information for ~15% of heterozygous SNP pairs from our genome sequencing data, we cannot rule out the possibility of underestimating the MNV mutation rate. As our sequencing technology as well as statistical and computational methods evolves, further analysis of this estimate would be valuable.

**(Analysis of distance d>2)**

In a similar way as we calculated the MNV density per genomic region, we calculated relative (overall) density of MNV against MNV of distance 10, for all the distance d=1..9 and all the MNV patterns. Specifically, the relative MNV density $D_r$ of distance d, pattern $W(*)_{d-1}X \rightarrow Y(*)_{d-1}Z$ , where $(*)_{d-1}$ denotes any sequence of length $(d-1)$, is defined as:

$$D_r\ (W, X \rightarrow Y, Z \mid d) = \frac{N(W(*)_{d-1}X \rightarrow Y(*)_{d-1}Z)\ /\ N(W(*)_{d-1}X)}{N(W(*)_9 X \rightarrow Y(*)_9 Z)\ /\ N(W(*)_9 X)} = \frac{N(W(*)_{d-1}X \rightarrow Y(*)_{d-1}Z)}{N(W(*)_9 X \rightarrow Y(*)_9 Z)} \cdot \frac{N(W(*)_9 X)}{N(W(*)_{d-1}X)}$$

According to this definition, $D_r\ (W, X \rightarrow Y, Z \mid d) = k$ means that the probability of observing a mutation of $W(*)_{d-1}X \rightarrow Y(*)_{d-1}Z$ given a sequence context of $W(*)_{d-1}X$ is k times higher than the probability of observing a mutation of $W(*)_9 X \rightarrow Y(*)_9 Z$ given a sequence context of $W(*)_9 X$. Also, the result of density calculation stratified by the functional annotation is available in Supplementary Fig. 20.
We did not expand our analysis to a range longer than 10 bp, because of the instability of read based phasing sensitivity and specificity (Supplementary Fig. 1). The base pattern in the supplementary figures denotes the reference and alternative 2 bp pattern, but does not specify the patterns of bases in between (Therefore, we do not exclude the possibility that an MNV of distance>2 is a subset of an MNV of larger window. For example, substitution of AAAA->TTTT would be counted as all of A,A->T,T of distance 1, 2, and 3).

### Supplementary Files legend

**Supplementary file 1. Definition of MNV functional category classification and counts in gnomAD exome, list of genes with more than zero gained nonsense mutation or rescued nonsense mutation, and the full list of MNV count per gene per category**

**Supplementary file 2. List of gained nonsense mutation, changed and gained missense with high CADD score, observed in rare disease samples, with a brief phenotype breakdown of the samples.**

**Supplementary file 3. Estimation of MNV frequency per MNV pattern per generation, and the breakdown of predicted major mechanism for each MNV pattern**

### Supplementary tables

**Table S1. Summary of phasing sensitivity experiment**

**a,** percentage for exome and genome. **b,c,** The raw number for exome **(b)** and genome**(c)**. **d,** The number and percentage of low and high depth variants, for each of the phased and unphased pairs. Thresholds: site depth $10^5$ for “low” and $10^6$ for “high” for genome ($10^6$and$10^{7.5}$for exome). Odds ratio =3.20, Fisher’s exact test $p < 10^{-100}$for low, and odds ratio =2.33, Fisher’s exact test $p < 10^{-100}$for high read depth in genome (2.19 and 1.51 for exome, $p < 10^{-100}$ and $p < 10^{-8}$). Chromosome 20 of randomly selected 10% of samples in gnomAD genome dataset, and all chromosome of 1% of samples in exome dataset was examined (we down sampled to roughly match the trio analysis in number while keeping the statistical power, and manually set the threshold by looking at the distribution).

**a. Genomes, 635 trios *( Exomes, 5785 trios )***

| Categ \ distance (bp) | 1 | 2 | .. | 10 | .. | 100 |
|---|---|---|---|---|---|---|
| % (both are phased) | 85.2 *(86.6)* | 76.3 *(84.7)* | .. | 80.6 *(81.5)* | .. | 2.30 *(0.412)* |
| %(has PBT) | 57.8 *(63.3)* | 53.9 *(57.2)* | .. | 51.4 *(55.0)* | .. | 17.0 *(26.1)* |
| %(agrees PBT) | 99.9 *(99.8)* | 99.5 *(99.6)* | .. | 99.4 *(99.1)* | .. | 91,5 *(61.5)* |

**b. (exome)**

| Categ \ distance (bp) | 1 | 2 | .. | 10 | .. | 20 | .. | 100 |
|---|---|---|---|---|---|---|---|---|
| all | 1245684 | 597400 | .. | 424871 | .. | 400862 | .. | 212742 |
| Same PID | 1078326 | 506155 | .. | 346068 | .. | 126485 | .. | 878 |
| Has PBT | 748658 | 326701 | .. | 215659 | .. | 179400 | .. | 104608 |
| MNV | 1023154 | 478736 | .. | 327943 | .. | 122539 | .. | 876 |

| | | | | | | | | |
|---|---|---|---|---|---|---|---|---|
| MNV that has PBT | 646261 | 270426 | .. | 177378 | .. | 50843 | .. | 229 |
| MNV that agrees PBT | 644761 | 270073 | .. | 177035 | .. | 50685 | .. | 125 |

**c. (genome)**

| Categ \ distance (bp) | 1 | 2 | .. | 10 | .. | 20 | .. | 100 |
|---|---|---|---|---|---|---|---|---|
| all | 15910983 | 10765039 | .. | 6790274 | .. | 5829142 | .. | 4754207 |
| Same PID | 13553340 | 8213791 | .. | 5473503 | .. | 2595977 | .. | 109656 |
| Has PBT | 8647090 | 5464182 | .. | 3224469 | .. | 2818533 | .. | 2371470 |
| MNV | 12996502 | 7891523 | .. | 5187226 | .. | 2484572 | .. | 108514 |
| MNV that has PBT | 7496987 | 4229588 | .. | 2634303 | .. | 1119211 | .. | 18471 |
| MNV that agrees PBT | 7487685 | 4221780 | .. | 2629642 | .. | 1116105 | .. | 17549 |

**d.**

| Categ \ depth bin | low | middle | high |
|---|---|---|---|
| number in genome, for unphased pair (%) | 796 (0.0308) | 2159905 (83.7) | 421327 (16.3) |
| number in genome, for phased pair (%) | 1129 (0.00962) | 10911850 (93.0) | 821621 (7.00) |

| | | | |
|---|---|---|---|
| number in exome, for unphased pair (%) | 292<br>(0.441) | 58358<br>(88.1) | 7564<br>(11.4) |
| number in exome, for phased pair (%) | 1090<br>(0.290) | 354753<br>(94.5) | 19581<br>(5.22) |

**Table S2. Description of the functional annotations used in the research**
The interval length, mean (across sites) of the median coverage (across individuals) in gnomAD, and the mean (across sites) methylation level of CpG sites are annotated as separate columns.

| Category | Interval length | % of genome | Coverage | Methylation level |
|---|---|---|---|---|
| TSS | 55509841 | 0.019603 | 30.6 | 0.194 |
| 5’ UTR | 19902177 | 0.007028 | 30.1 | 0.350 |
| Promoter | 94046133 | 0.033212 | 30.4 | 0.437 |
| Enhancer | 126128190 | 0.044541 | 31.1 | 0.448 |
| TFBS | 380345514 | 0.134316 | 31.2 | 0.502 |
| H3K4me3 | 397134049 | 0.140244 | 31.3 | 0.532 |
| DHS | 492285933 | 0.173846 | 31.3 | 0.543 |
| H3K9ac | 376617506 | 0.132999 | 31.2 | 0.546 |
| Coding | 61730033 | 0.021799 | 31.2 | 0.584 |
| H3K27ac | 772035773 | 0.272638 | 30.9 | 0.664 |
| H3K4me1 | 1242793221 | 0.438882 | 31.1 | 0.717 |
| 3’ UTR | 40715655 | 0.014378 | 30.7 | 0.720 |
| Intron | 1123766231 | 0.396848 | 30.6 | 0.797 |
| Transcribed | 1022244971 | 0.360997 | 30.4 | 0.866 |

**Table S3. Summary of the study**

RF stands for random forest filtering, and adj stands for adjusted threshold (GQ >= 20, DP >= 10, and have now added: allele balance > 0.2 for heterozygote genotypes) filtering for each sample. 129 variants in Rare disease exome had gained nonsense mutation or high CADD score and low gnomAD frequency, but none of them were likely to be causal after manual inspection.

| Callset | Exome | Genome | Rare disease exome |
|---|---|---|---|
| Sample size | 125,748 | 15,708 | 6,072 |
| Filtering Criteria | RF / adj | RF | N/A |
| Annotations | - amino acid change<br>- gene constraint | - sequence context<br>- methylation status<br>- functional category | - gene constraint<br>- CADD score<br>- frequency in gnomAD |
| Number of MNVs | 31,510 (within codon) | 1,996,125 | 129 (Not likely causal) |
| Main usage | Functional impact analysis | Mutational mechanisms analysis | Clinical diagnosis |

**Table S4. Table view of the validation of phasing accuracy of read-based phasing using trio-based phase information**

We are showing the case where read based phase of SNP1 is 0|1, without loss of generality.

<table>
<tr><th>SNP1<br>SNP2<br>(read base phase)</th><th>SNP1<br>SNP2<br>(trio base phase)</th><th>consistent</th></tr>
<tr><td>0|1<br>0|1</td><td>0|1<br>0|1</td><td>True</td></tr>
<tr><td>0|1<br>0|1</td><td>0|1<br>1|0</td><td>False</td></tr>
<tr><td>0|1<br>0|1</td><td>1|0<br>0|1</td><td>False</td></tr>
<tr><td>0|1<br>0|1</td><td>1|0<br>1|0</td><td>True</td></tr>
<tr><td>0|1<br>1|0</td><td>0|1<br>0|1</td><td>False</td></tr>
<tr><td>0|1<br>1|0</td><td>0|1<br>1|0</td><td>True</td></tr>
</table>

| | | |
|---|---|---|
| 0\|1<br>1\|0 | 1\|0<br>0\|1 | True |
| 0\|1<br>1\|0 | 1\|0<br>1\|0 | False |

**Supplementary figures:**

**a.**

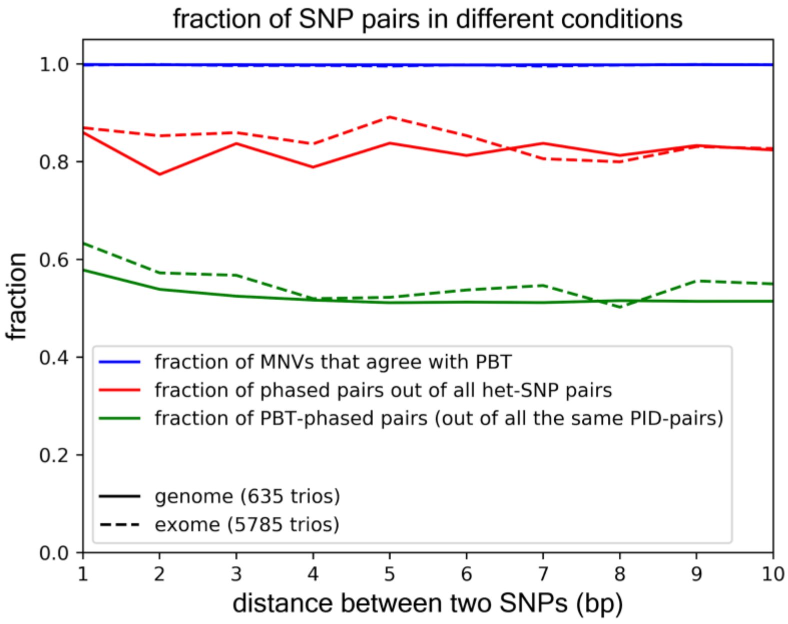

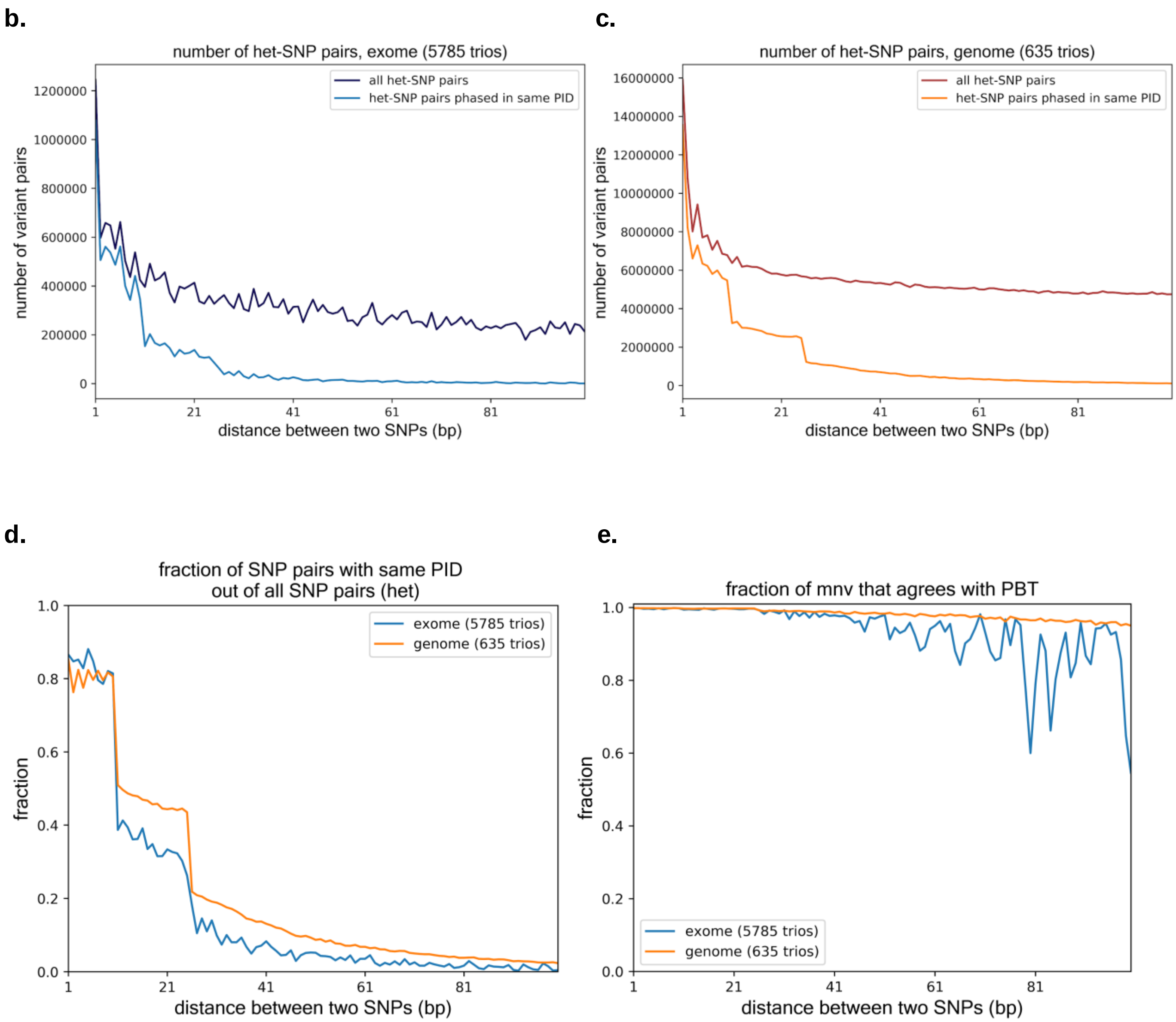


**Figure S1. Phasing quality as a function of distance**

**a,** Fraction of MNVs that agrees with PBT (trio based phasing information), fraction of phased heterozygous SNP pairs that has phase information (by read base phasing), and fraction of trios that has trio based phasing information, up to 10 bp. Trio based phasing is by definition impossible when both of the parents are heterozygous, resulting in relatively low phasing sensitivity. **b-d,** The number **(b,c)** and the fraction **(d)** of heterozygous SNP pairs that has same Phase ID, out of all the heterozygous SNP pairs, up to 100 bp. **e,** Fraction of MNVs that agrees with trio based phase information, up to 100 bp.

For **(b)~(e)**, the estimation becomes unstable as the distance becomes large, because of both the limited number of sample size and phasing sensitivity.

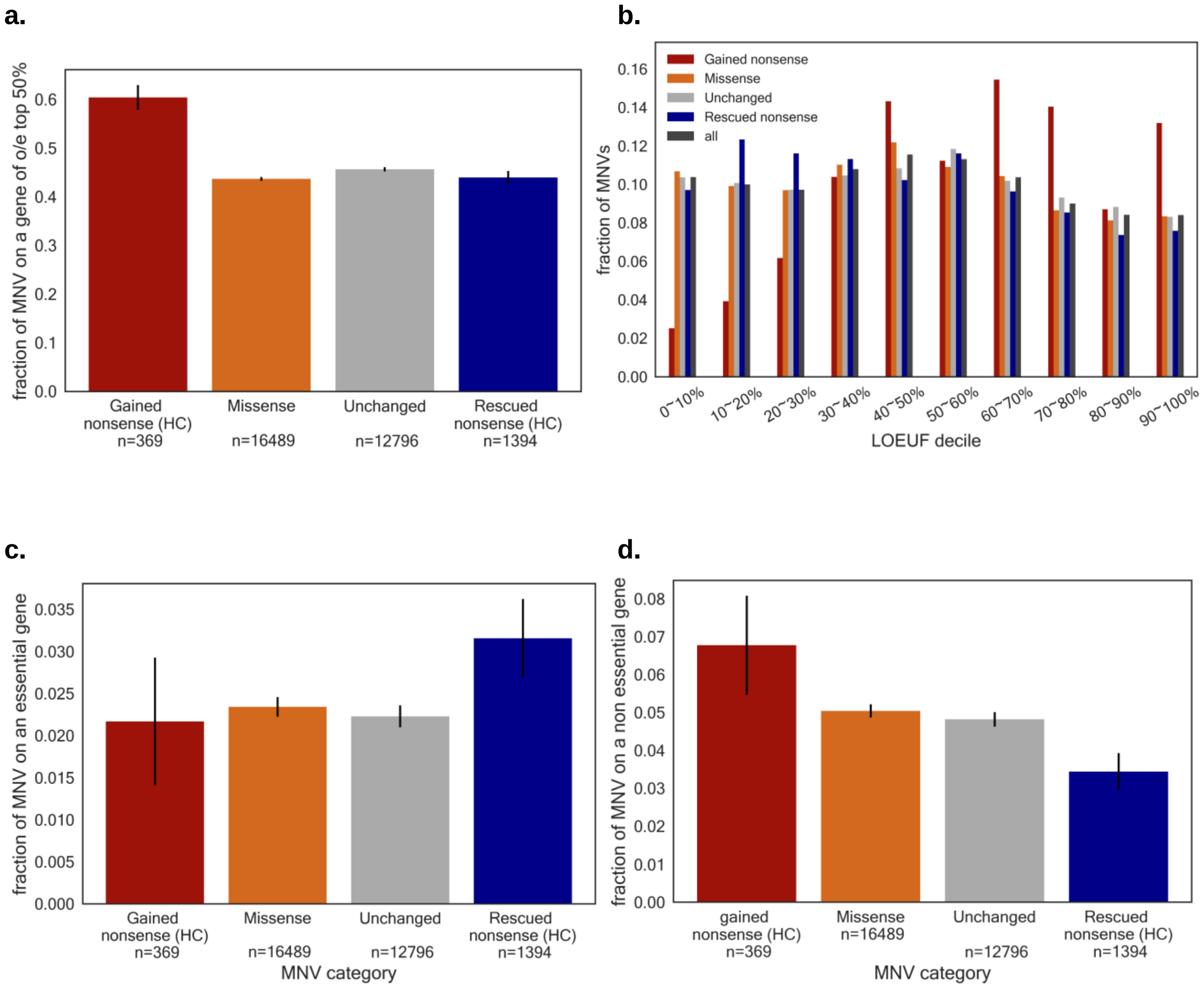


**Figure S2. MNV and gene constraint, cell essentiality**

**a,** Fraction of MNV that falls into un-constrained genes (n=9852) defined as top 50% in the observed vs expected number of loss-of-function in gnomAD data (LOEUF>0.926). The fraction for gained nonsense mutation is significantly higher than all the other classes combined (Fisher's exact test p-value < $7.04 \times 10^{-5}$). **b,** Fraction of MNVs of each functional category represented in each LOEUF decile. X axis shows the LEOUF decile, and y axis is fraction of MNVs that fall into genes of that specific LEOUF decile. **c,** Fraction of MNV that falls into essential genes (n=684), defined by knockout screening experiments using CRISPR/Cas[43,44]. The fraction for rescued nonsense mutation is significantly higher than all the other classes combined (Fisher's exact test p-value < 0.047). **d,** Fraction of MNV that falls into non essential genes (n=928), defined by knockout screening experiments using CRISPR/Cas[43,44] . The fraction for rescued nonsense mutation is significantly lower than all the other classes combined (Fisher's exact test p-value < 0.011).

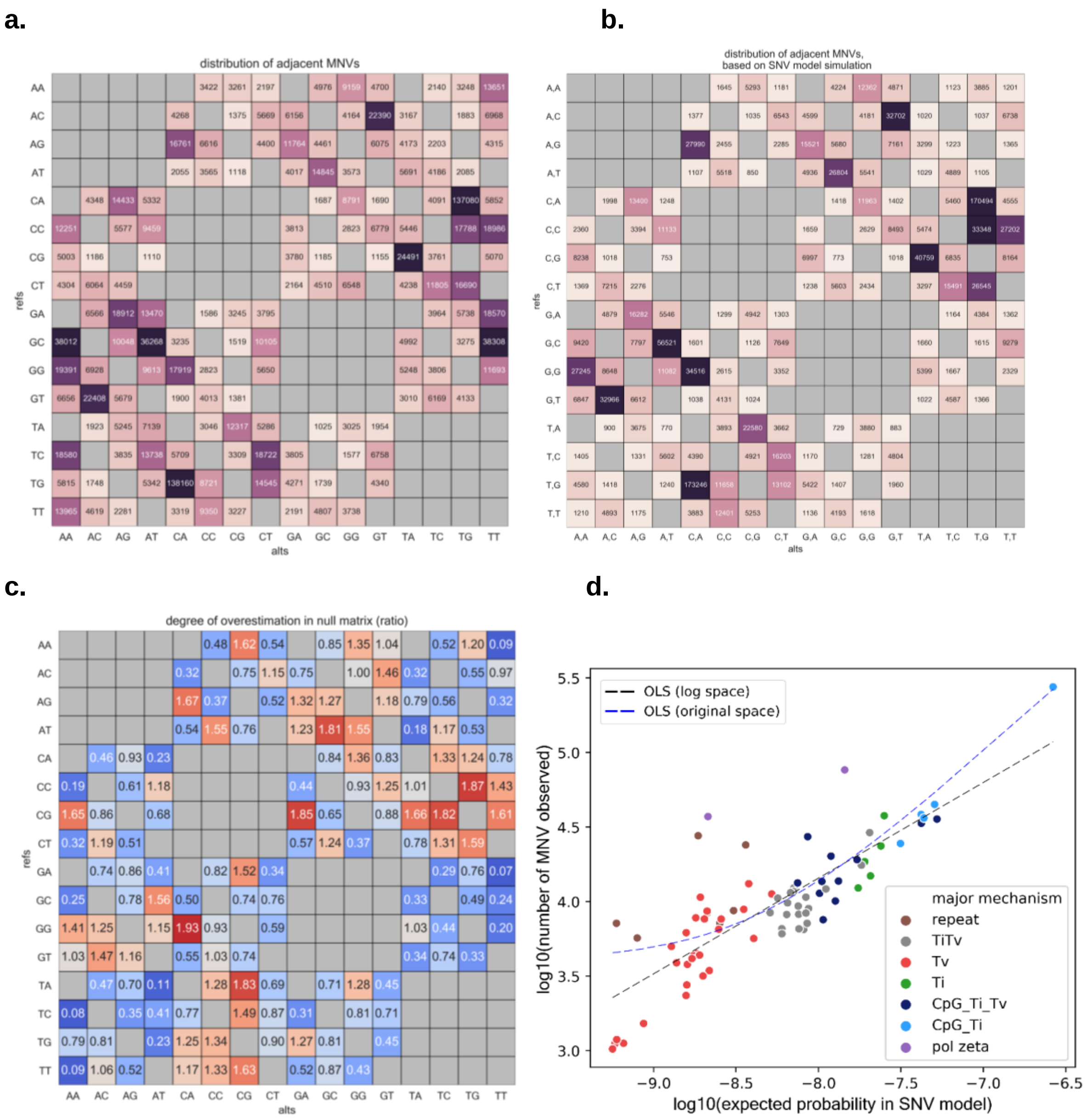


**Figure S3. Matrix representation of number of adjacent MNVs, and comparison with simulation by simple probabilistic model**

**a,** Matrix representation of number of adjacent MNVs (reverse complements are not collapsed). Grey colored ones are by definition not MNV (SNP or not even a mutation). **b,** Matrix representation of number of adjacent MNVs (reverse complements are not collapsed), in a null model based on SNV mutation rate, , assuming all MNV are consequence of two single nucleotide variation events. **c,** Enrichment compared to the null model based on SNV mutation rate. **d,** Scatter plot where x axis shows the relative probability of MNV based on null model (per generation), and y axis is the number of MNV observed in gnomAD data, both in log10 scale. The color represents the predicted major mechanism for each MNV pattern. Polymerase zeta and repetitive motifs are pushing up the number of MNV compared to the null model. Pearson correlation r=0.82 before removing polymerase zeta and repeat signature, and r=0.99 after removing those two signature (purple and brown), for linear regression in log space (black line).

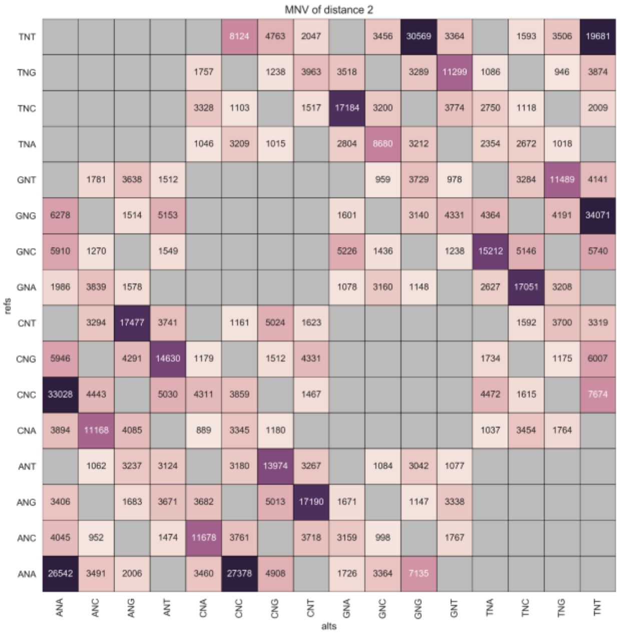
MNV of distance 2
refs
alts
TNT
TNG
TNC
TNA
GNT
GNG
GNC
GNA
CNT
CNG
CNC
CNA
ANT
ANG
ANC
ANA

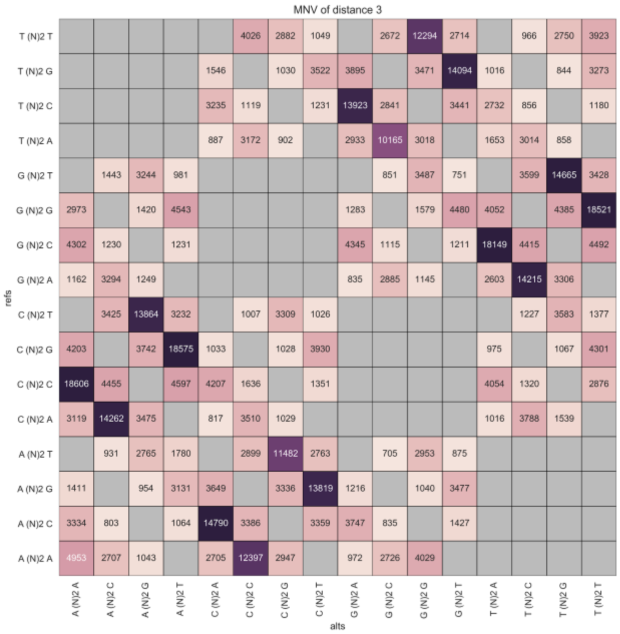
MNV of distance 3
refs
alts
T (N)2 T
T (N)2 G
T (N)2 C
T (N)2 A
G (N)2 T
G (N)2 G
G (N)2 C
G (N)2 A
C (N)2 T
C (N)2 G
C (N)2 C
C (N)2 A
A (N)2 T
A (N)2 G
A (N)2 C
A (N)2 A

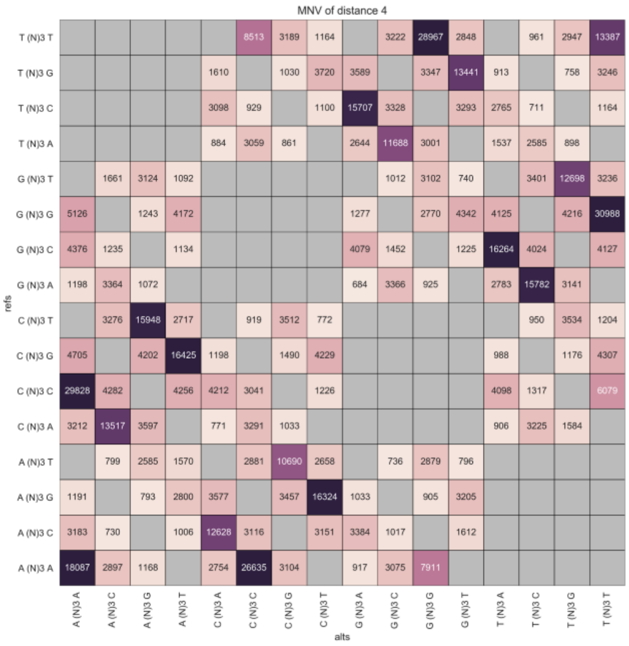
MNV of distance 4
refs
alts
T (N)3 T
T (N)3 G
T (N)3 C
T (N)3 A
G (N)3 T
G (N)3 G
G (N)3 C
G (N)3 A
C (N)3 T
C (N)3 G
C (N)3 C
C (N)3 A
A (N)3 T
A (N)3 G
A (N)3 C
A (N)3 A

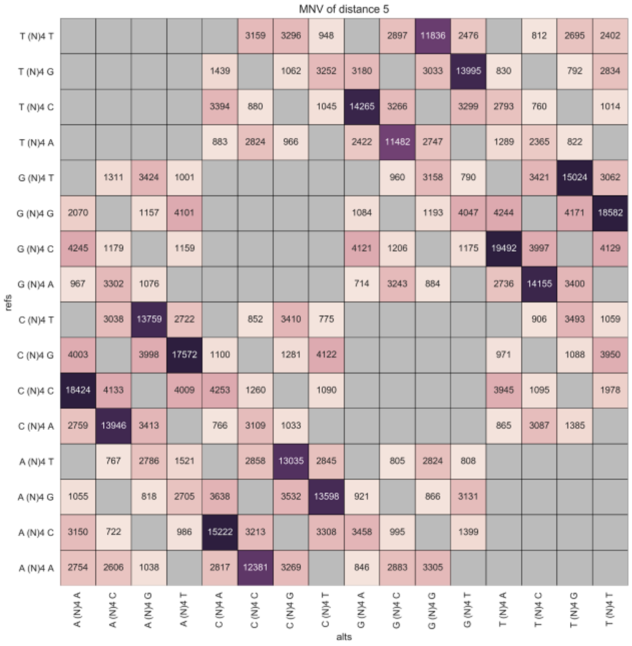
MNV of distance 5
refs
alts
T (N)4 T
T (N)4 G
T (N)4 C
T (N)4 A
G (N)4 T
G (N)4 G
G (N)4 C
G (N)4 A
C (N)4 T
C (N)4 G
C (N)4 C
C (N)4 A
A (N)4 T
A (N)4 G
A (N)4 C
A (N)4 A

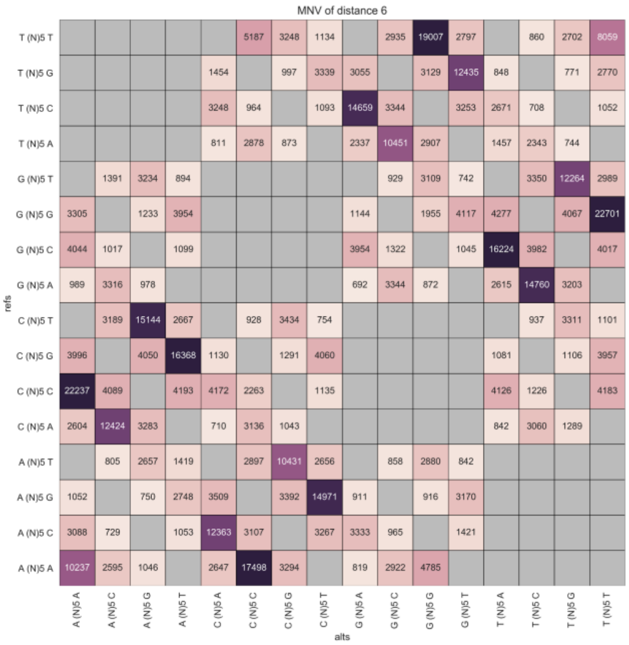
MNV of distance 6
refs
alts

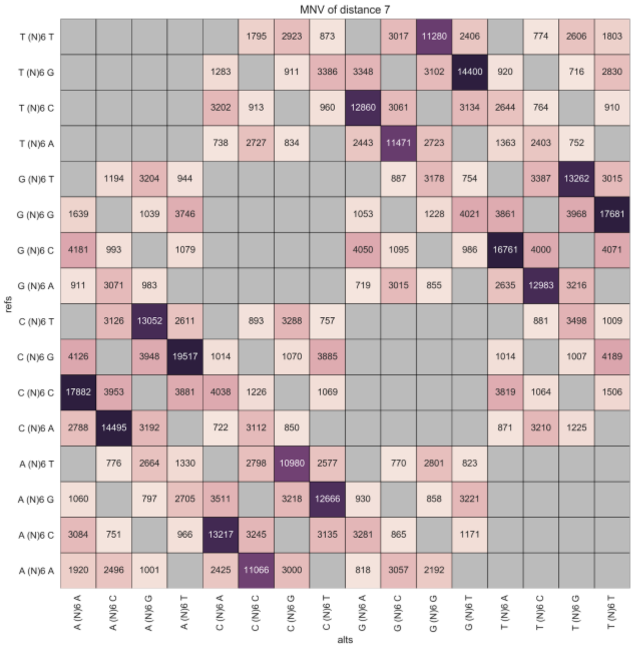
MNV of distance 7
refs
alts

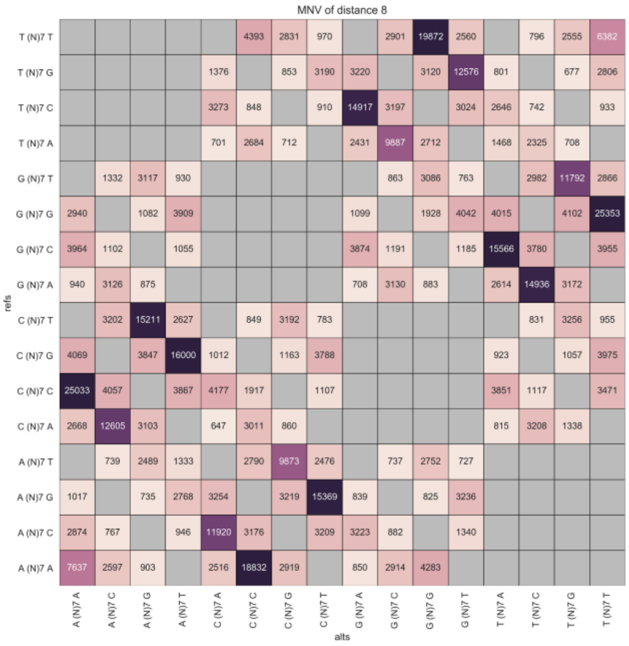
MNV of distance 8
refs
alts

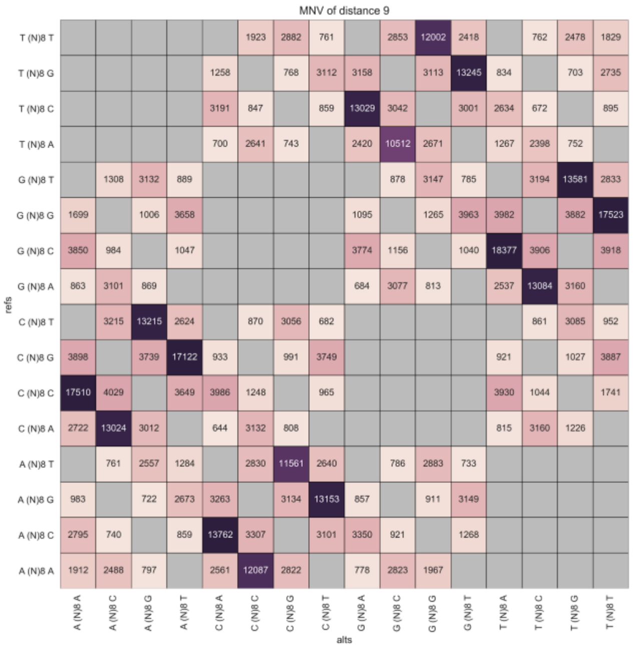
MNV of distance 9
refs
alts

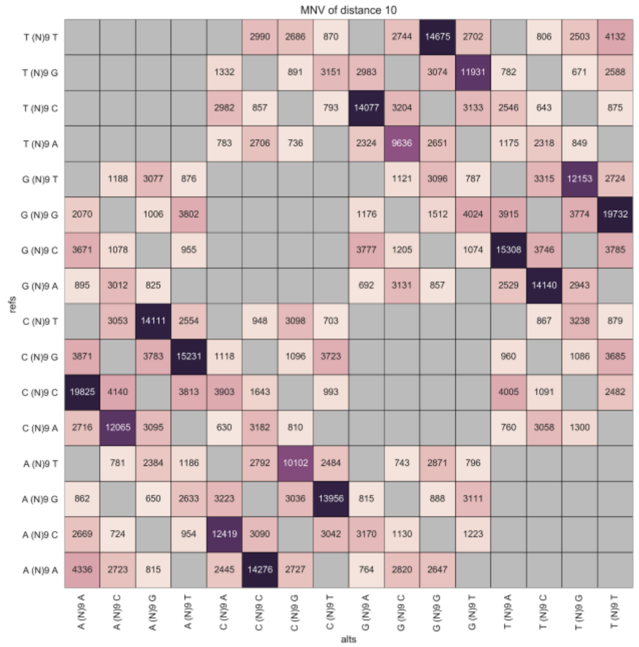


**Figure S4. Matrix representation of number of MNVs, up to 10 bp**

Row denotes the reference, and the column denotes the alternative alleles. (N)M in the row and column means arbitral sequence of length M (e.g. (N)2 is one of {AA,AC,AG,...,TT}).

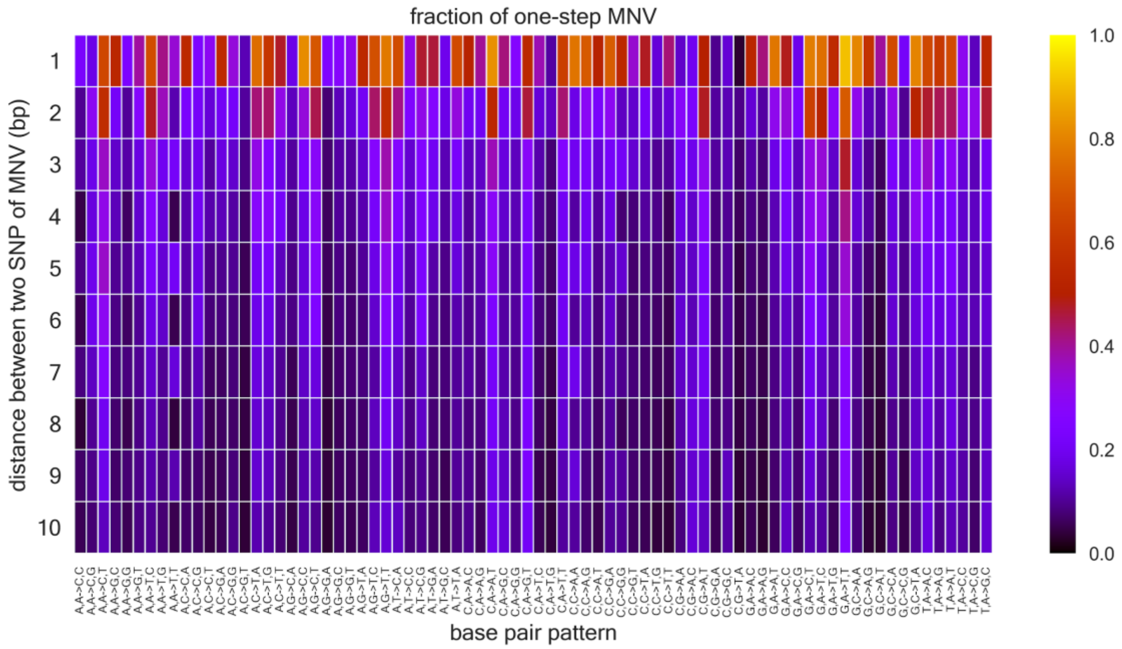


**Figure S5. Fraction of one-step MNV, up to 10 bp**

Row denotes the distance between two SNPs of MNV, and column denotes the base pair pattern. The fraction is represented as color, although the color scale is different for each distance.

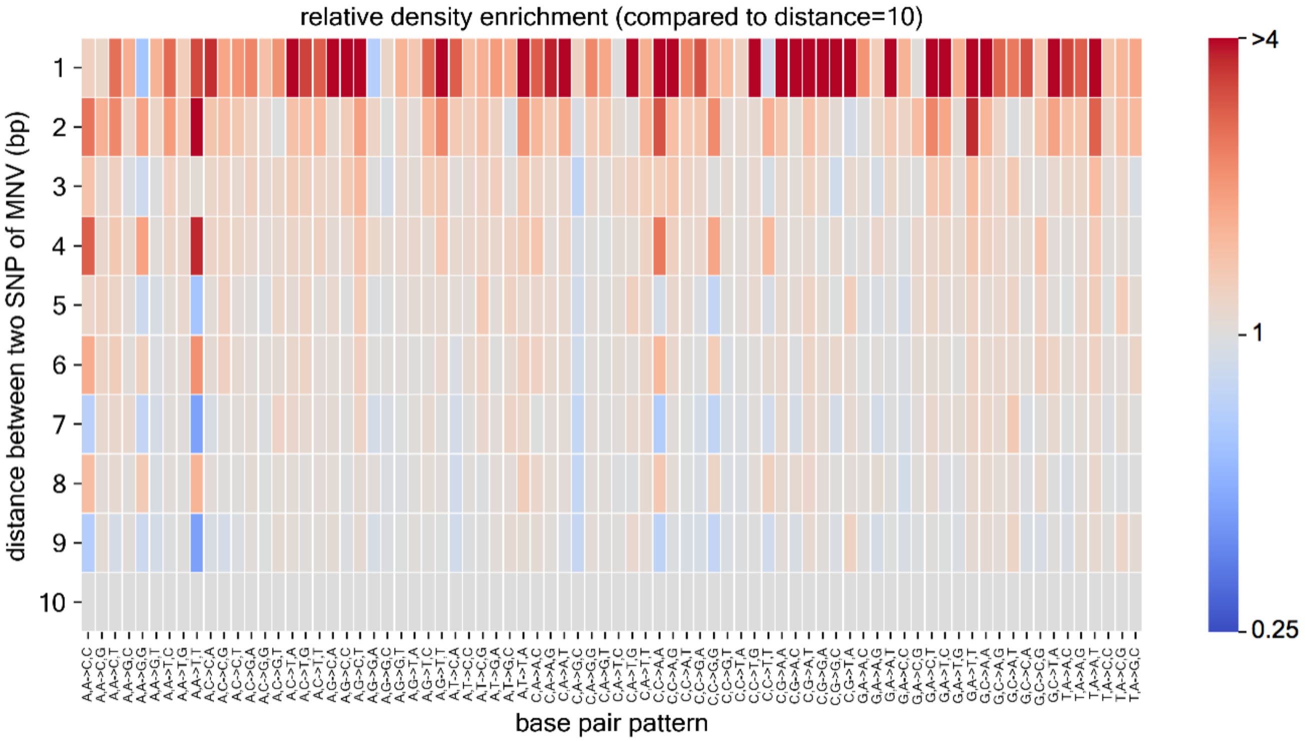


**Figure S6. Relative MNV density enrichment per MNV pattern, compared to 10 bp distance**
Row denotes the distance between two SNPs of variant pairs, and column denotes the base pair pattern. The relative density enrichment is represented as color (color shifts in log scale). By definition for row=10, all the column entries are exactly equal to 1.

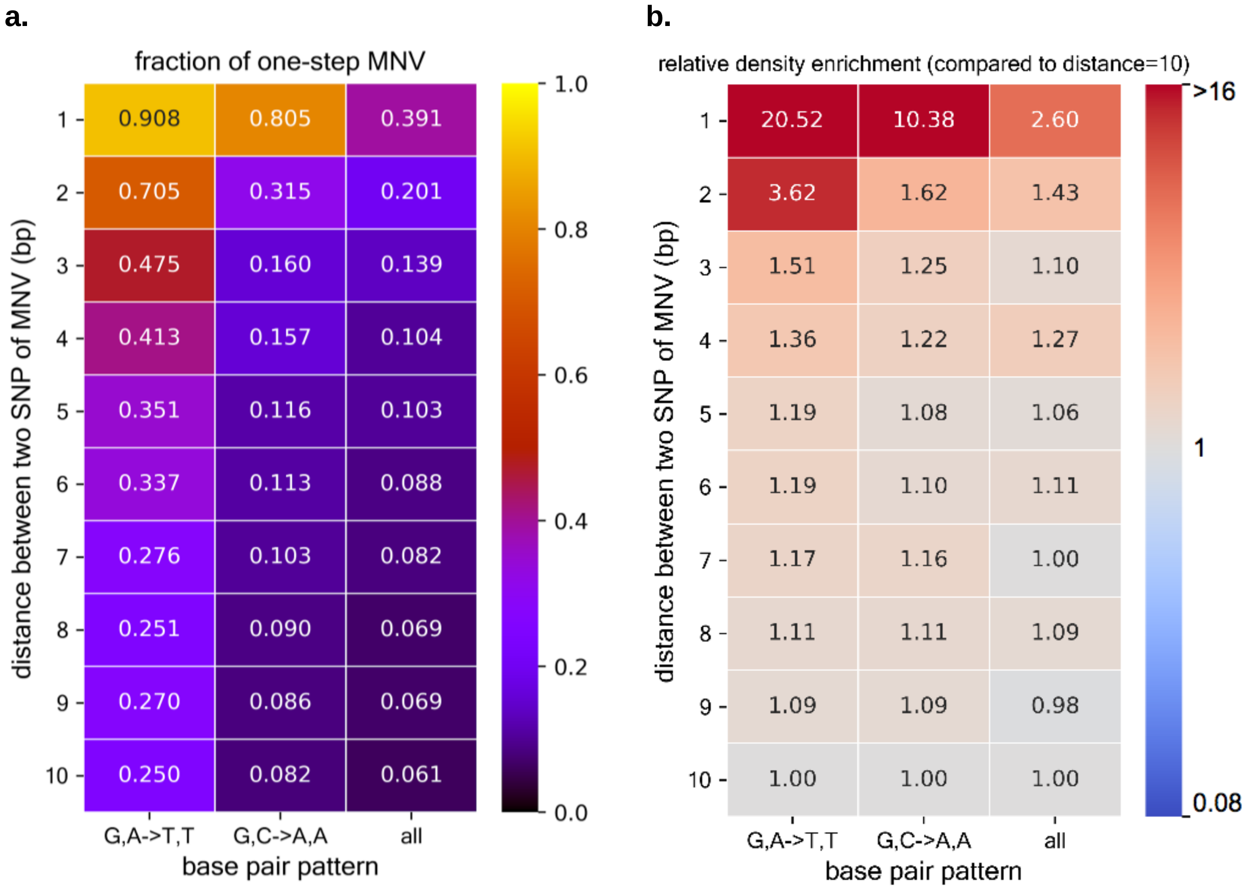


| distance (bp) | a. G,A->T,T | a. G,C->A,A | a. all | b. G,A->T,T | b. G,C->A,A | b. all |
|---|---|---|---|---|---|---|
| 1 | 0.908 | 0.805 | 0.391 | 20.52 | 10.38 | 2.60 |
| 2 | 0.705 | 0.315 | 0.201 | 3.62 | 1.62 | 1.43 |
| 3 | 0.475 | 0.160 | 0.139 | 1.51 | 1.25 | 1.10 |
| 4 | 0.413 | 0.157 | 0.104 | 1.36 | 1.22 | 1.27 |
| 5 | 0.351 | 0.116 | 0.103 | 1.19 | 1.08 | 1.06 |
| 6 | 0.337 | 0.113 | 0.088 | 1.19 | 1.10 | 1.11 |
| 7 | 0.276 | 0.103 | 0.082 | 1.17 | 1.16 | 1.00 |
| 8 | 0.251 | 0.090 | 0.069 | 1.11 | 1.11 | 1.09 |
| 9 | 0.270 | 0.086 | 0.069 | 1.09 | 1.09 | 0.98 |
| 10 | 0.250 | 0.082 | 0.061 | 1.00 | 1.00 | 1.00 |

**Figure S7. Polymerase zeta signatures are observed for distance>1**
Figure generated by taking two columns from **figure S5** and **S6**, and adding the overall level as comparison. The fraction of one-step MNV **(a)** and the density of MNV **(b)** for G,A->T,T and G,C->A,A, the MNV pattern known as polymerase zeta signature when the distance is 1 bp, are higher compared to the overall value, even for distance>1 (= not only GA->TT, but also GNA->TNT, GNNA->TNNT, … are enriched) (Fisher's exact test p-value < 0.05 up to distance 6 bp for the fraction of one-step MNV, and up to 9 bp for the relative density, compared to MNVs of same pattern, distance 10 bp). We do not fully exclude the possibility that the signals we observe are driven by sequence error or other artifact, since the fraction of variants that are filtered out tends to be high and the mean coverage is low for these variants, especially when two base pairs of MNV are not adjacent.

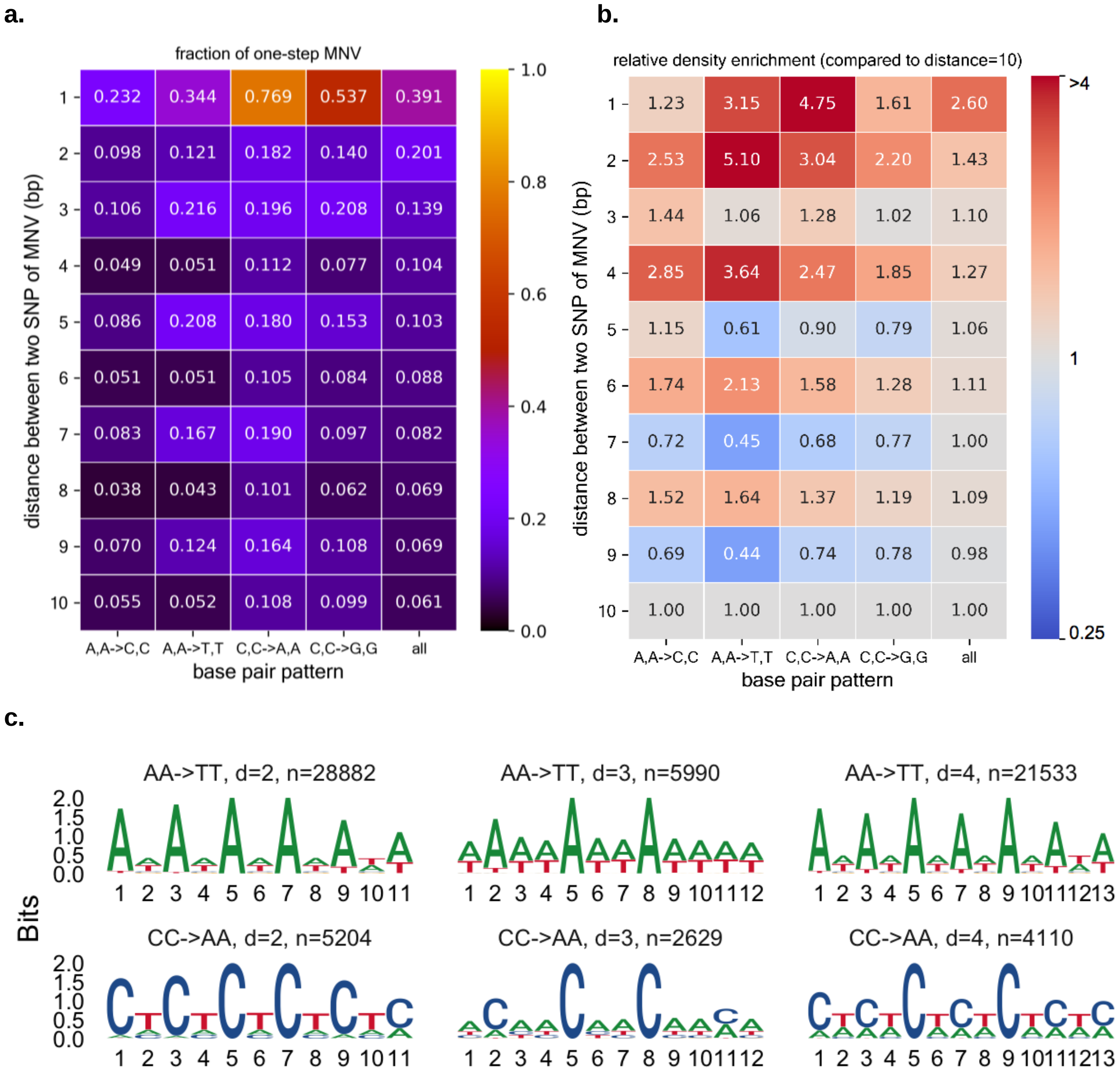
a.
fraction of one-step MNV
distance between two SNP of MNV (bp)
base pair pattern
A,A->C,C A,A->T,T C,C->A,A C,C->G,G all
1 0.232 0.344 0.769 0.537 0.391
2 0.098 0.121 0.182 0.140 0.201
3 0.106 0.216 0.196 0.208 0.139
4 0.049 0.051 0.112 0.077 0.104
5 0.086 0.208 0.180 0.153 0.103
6 0.051 0.051 0.105 0.084 0.088
7 0.083 0.167 0.190 0.097 0.082
8 0.038 0.043 0.101 0.062 0.069
9 0.070 0.124 0.164 0.108 0.069
10 0.055 0.052 0.108 0.099 0.061
1.0 0.8 0.6 0.4 0.2 0.0
b.
relative density enrichment (compared to distance=10)
distance between two SNP of MNV (bp)
base pair pattern
A,A->C,C A,A->T,T C,C->A,A C,C->G,G all
1 1.23 3.15 4.75 1.61 2.60
2 2.53 5.10 3.04 2.20 1.43
3 1.44 1.06 1.28 1.02 1.10
4 2.85 3.64 2.47 1.85 1.27
5 1.15 0.61 0.90 0.79 1.06
6 1.74 2.13 1.58 1.28 1.11
7 0.72 0.45 0.68 0.77 1.00
8 1.52 1.64 1.37 1.19 1.09
9 0.69 0.44 0.74 0.78 0.98
10 1.00 1.00 1.00 1.00 1.00
>4 1 0.25
c.
Bits
AA->TT, d=2, n=28882
AA->TT, d=3, n=5990
AA->TT, d=4, n=21533
CC->AA, d=2, n=5204
CC->AA, d=3, n=2629
CC->AA, d=4, n=4110
2.0 1.5 1.0 0.5 0.0

**d.**

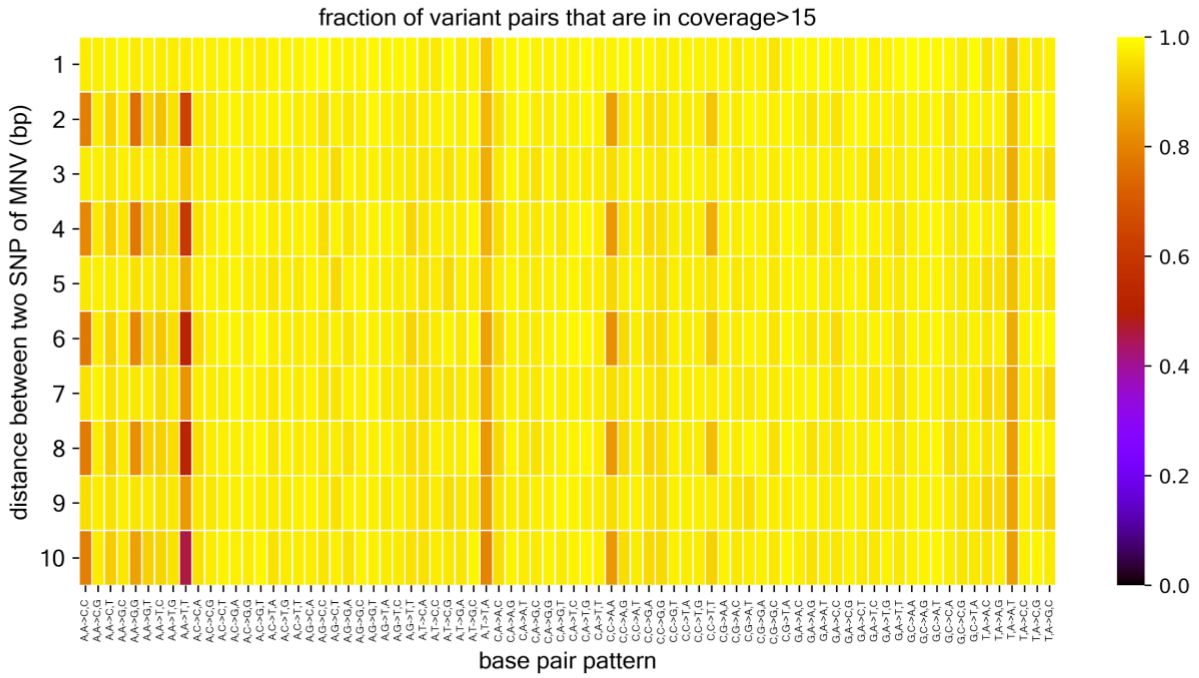


**Figure S8. Repeat signature are observed for distance>1, but has low coverage overall**
**a,** Figure generated by taking four columns from **figure S5** and **S6**, and adding the overall level as comparison. The fraction of one-step MNV **(a)** and the density of MNVature when the distance is 1 bp, are higher compared to the overall value, even for distance>1 (Fisher's exact test p-value < 0.05 up to distance 3 bp for the fraction of one-step MNV, and up to 9 bp for the relative density, compared to MNVs of same pattern, distance 10 bp), and favors distance of even number (e.g. ANNNA->TNNNT is enriched compared to ANNA->TNNT), presumably due to the instability of tandem repeat of 2 bp length unit. We do not fully exclude the possibility that the signals we observe are driven by sequence error or other artifact, for same reason as **figure S7,** and as shown in **(d)**. **c,** examples of enriched repetitive sequence context in distance 2, 3 and 4 (x=relative base pair position). **d,** Fraction of MNV that are in a region of median coverage >15, up to 10 bp. Row denotes the distance between two SNPs of variant pairs, and column denotes the MNV pattern (and reverse complement). The relative density enrichment is represented as color.

a. b.

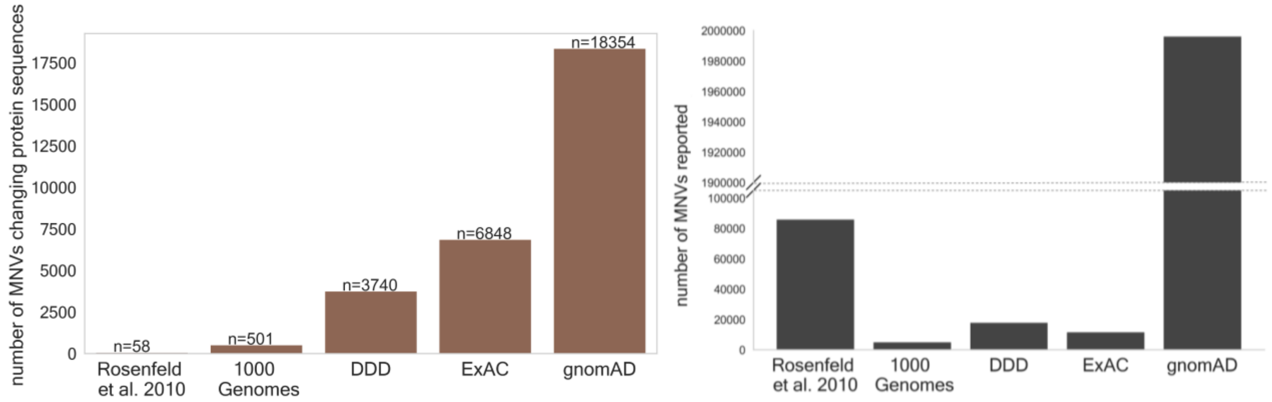


**Figure S9. Comparison of numbers of MNVs in different studies**

Number of MNVs that change protein sequences **(a)** and the total number of MNVs reported **(b)**, for five major studies to date: Rosenfeld et al. (2010)[3], 1000 Genomes[9,10], Deciphering Developmental Disorder (DDD)[2], Exome Aggregation Consortium (ExAC)[1], and genome Aggregation Database (gnomAD)[22]. For Rosenfeld et al. in **(a)**, we estimated the number from the total number of MNVs in exon, using the statistics from DDD and gnomAD. Note that **(b)** is not necessarily equal to the total number of MNVs discovered in the study, since some studies do not explicitly report non-coding MNVs. Also the statistics is subject to different variant calling and filtering criteria applied for different studies.

a. b.

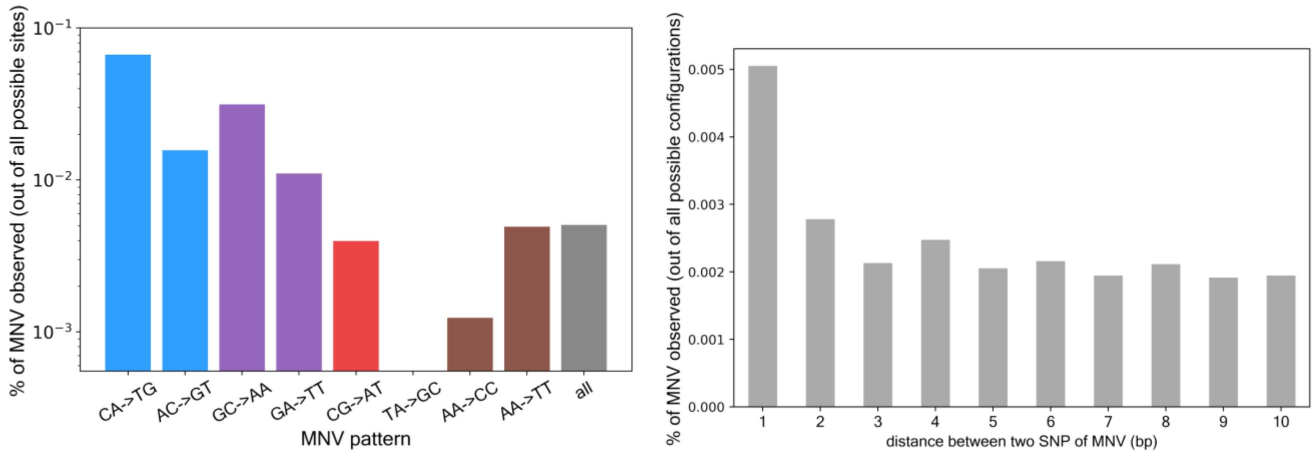


**Figure S10. percentage of MNV we have observed**

**a,** Percentage of MNV we have observed, per MNV pattern. Two patterns per major mutational mechanisms are shown as representations. Y axis is log scaled. 0.000551% for TA->GC. **b,** Percentage of MNV we have observed, per distance.

**a.**

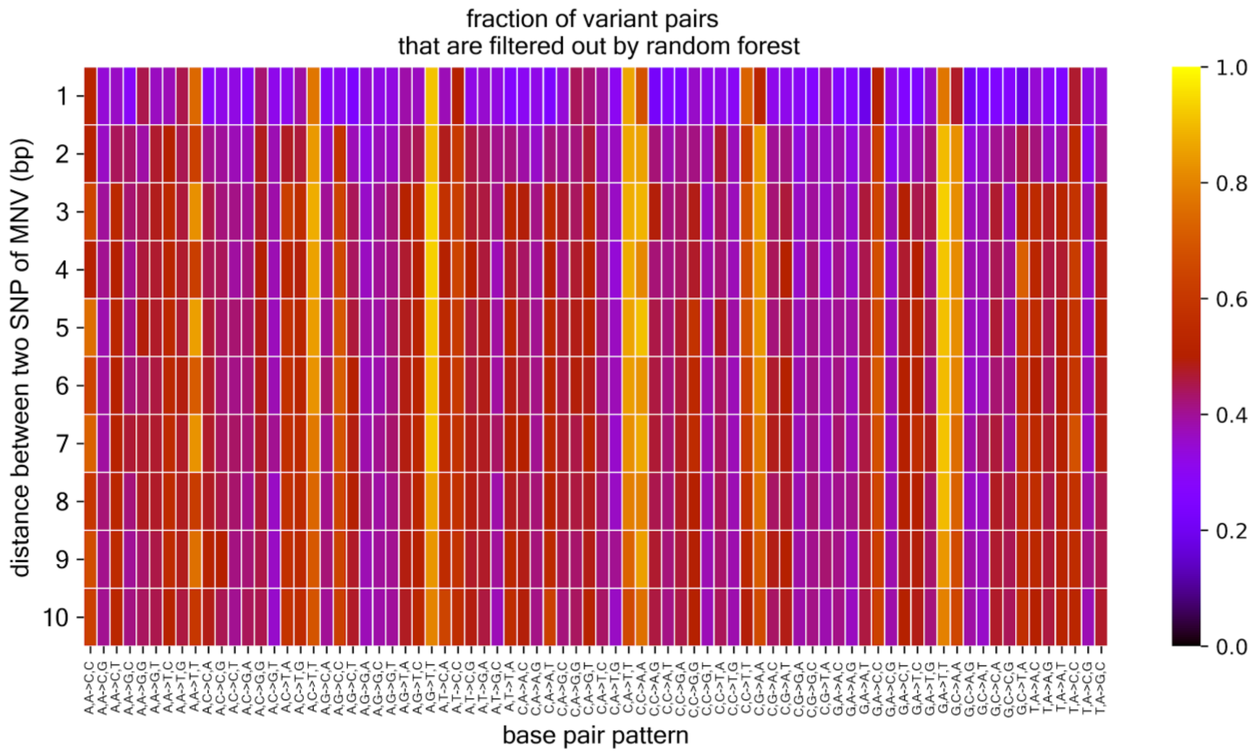


**b.**

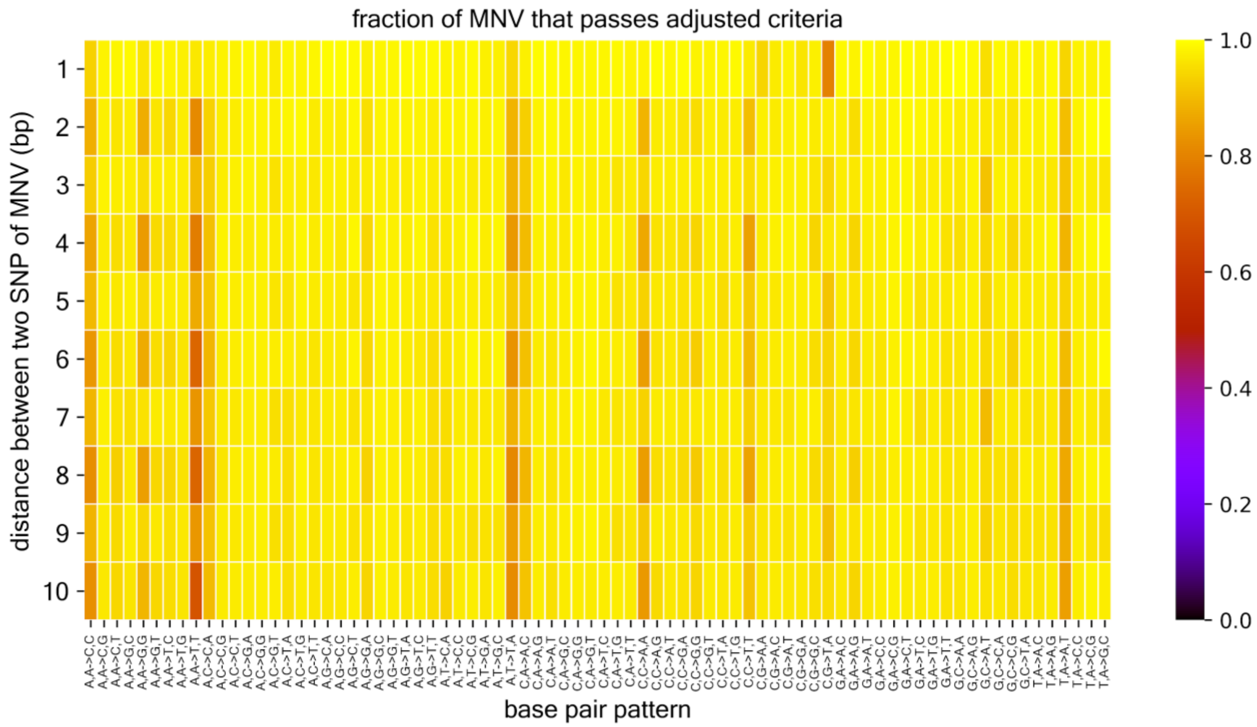


**Figure S11. Fraction of variant pairs that are filtered out by random forest (a) / passed the adjusted criteria (b), up to 10 bp**

Row denotes the distance between two SNPs of variant pairs, and column denotes the MNV pattern (and reverse complement). The fraction is represented as color. Since the fraction that pass adj criteria is high (>90% for >70 out of 78 patterns for distance 2 bp, >60% for all patterns across all distance, and >80% overall), we did not apply adj criteria for genome analysis, to obtain larger number of MNVs.

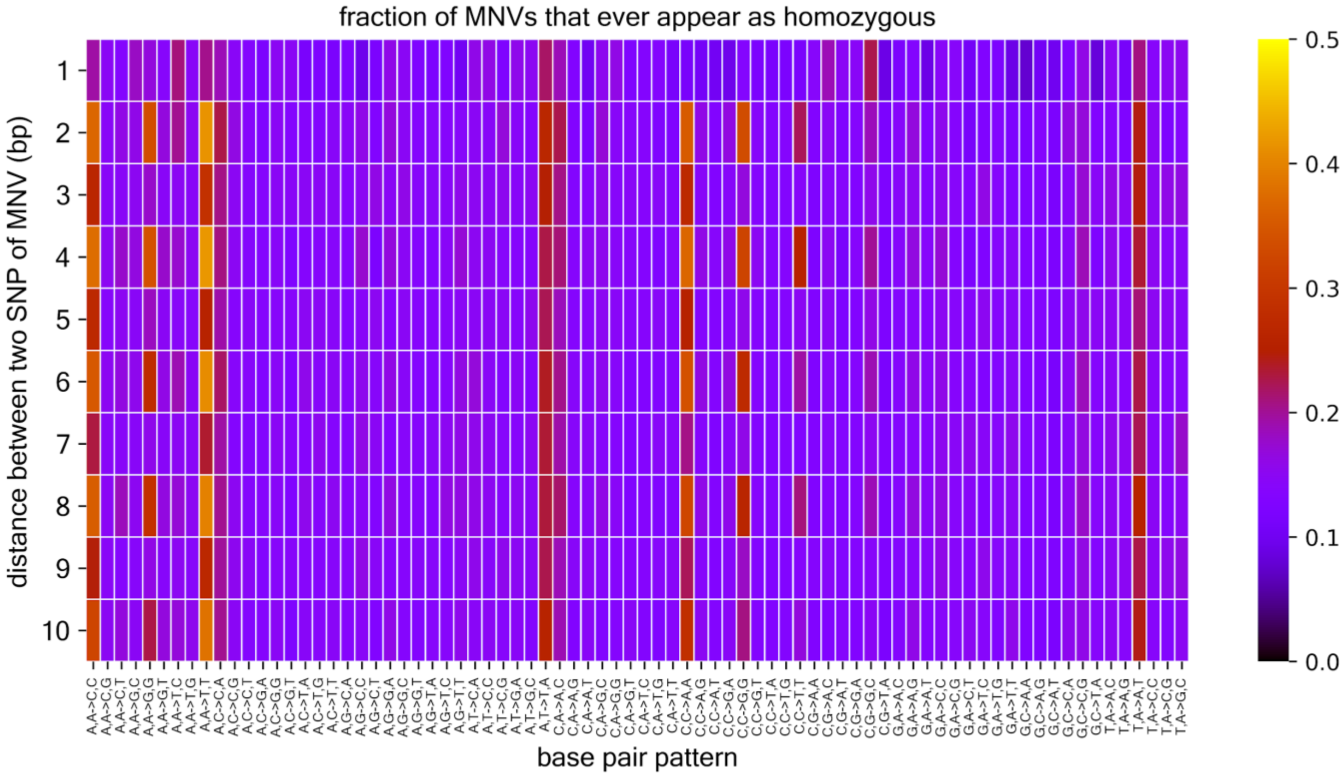


**Figure S12. Fraction of MNV that homozygous MNV are observed, up to 10 bp**
Row denotes the distance between two SNPs of variant pairs, and column denotes the MNV pattern. The relative density enrichment is represented as color.

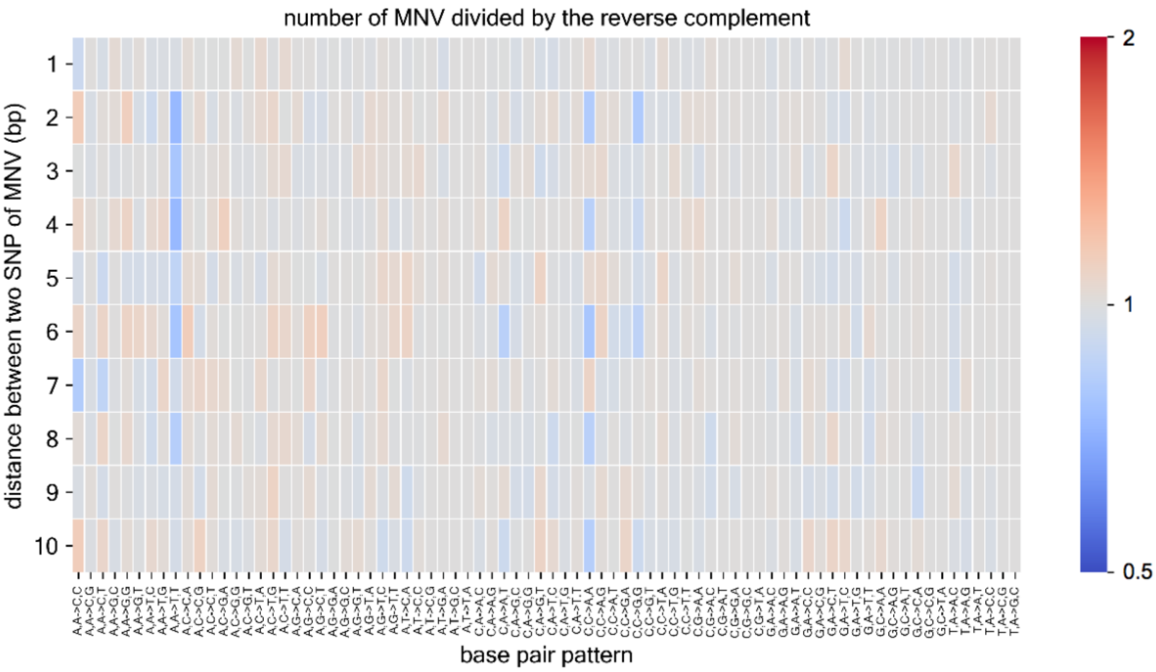


**Figure S13. ratio of number of MNV against their reverse complements, up to 10 bp**
Row denotes the distance between two SNPs of variant pairs, and column denotes the MNV pattern. The ratio against their reverse complement is represented as color (color shifts in log scale). Except for this figure, all the MNV pattern (d=1), and base pair pattern (d=1..10) do not distinguish the corresponding reverse complements.

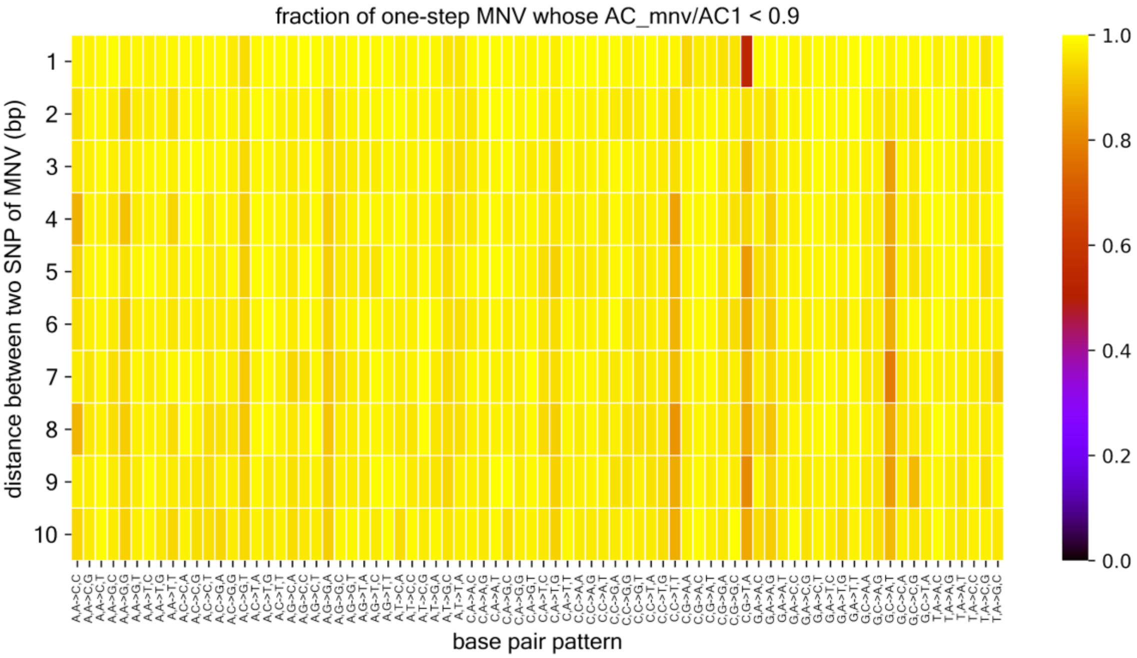


**Figure S14. Fraction of one-step MNV whose AC_mnv is close enough, up to 10 bp**

Row denotes the distance between two SNPs of variant pairs, and column denotes the MNV pattern. The fraction is represented as color. We took the threshold of AC_mnv / AC_1 >0.9 manually, after observing figure S7.

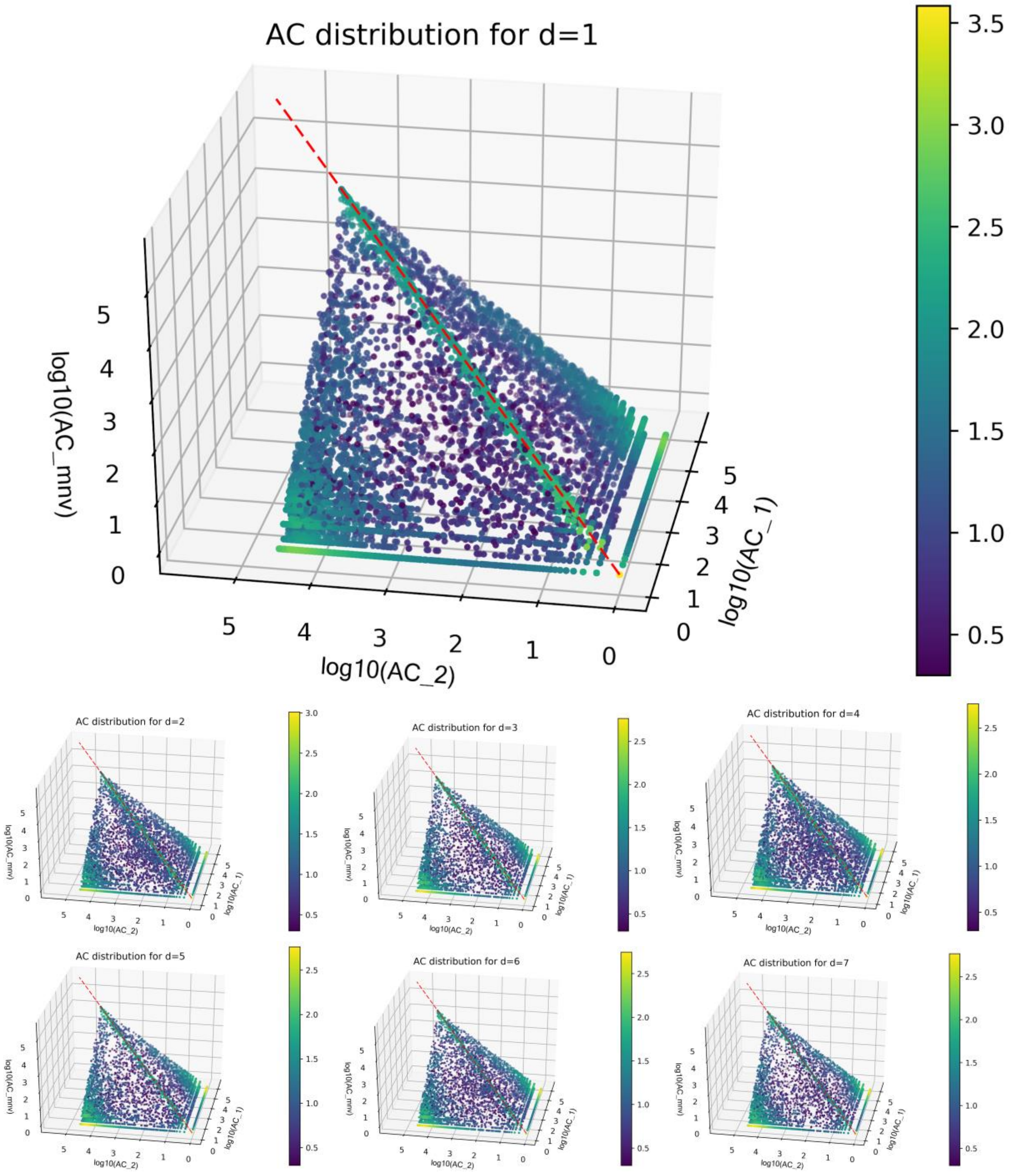

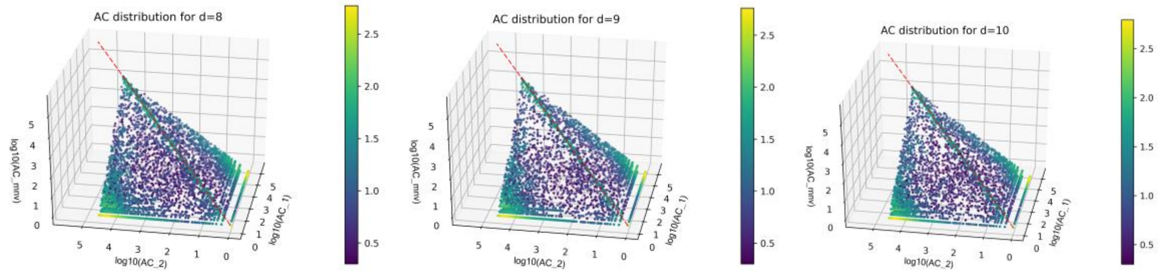


**Figure S15. Distribution of allele counts of MNVs, up to 10 bp**

One axis is the allele count of SNP that are upstream (AC_1), another for the allele count of SNP that are downstream (=AC_2) in the reference genome, and the other axis is for the allele count of corresponding MNV (in log10 space). Color corresponds to the relative density (defined as the number of neighborhood counts in log-space), and the read dot line shows x=y=z (i.e. AC of SNP1 = AC of SNP2 = AC of MNV). Only the result of chr22 is shown, in order for the figure to be sparse enough for effective visualization.

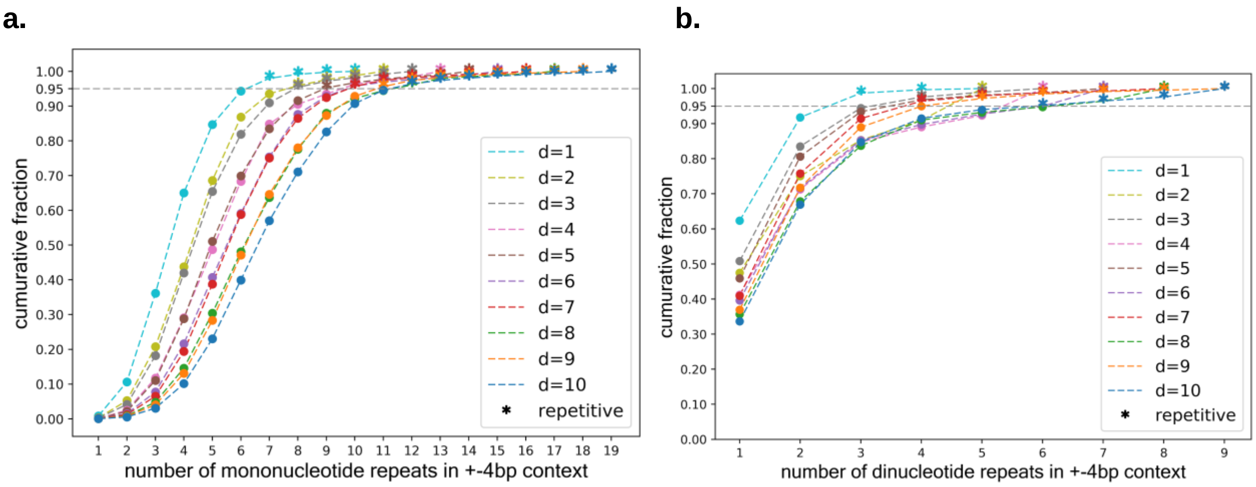


**Figure S16. Distribution of number of repeats**

**a, b,** x axis is the maximum number of mononucleotide **(a)** or dinucleotide **(b)** repeats (minimum of the reference +-4 bp and alternative +-4 bp context), and y is the cumulative density. Each line corresponds to specific distance between two SNP of MNV, and the start shape shows that they pass the threshold of cumulative percentage >95%, and thus are defined as “repetitive”. **c,** Fraction of MNV in repetitive contexts, defined as in **(a)** and **(b)**. Row denotes the distance between two SNPs of MNV, and column denotes the MNV pattern.

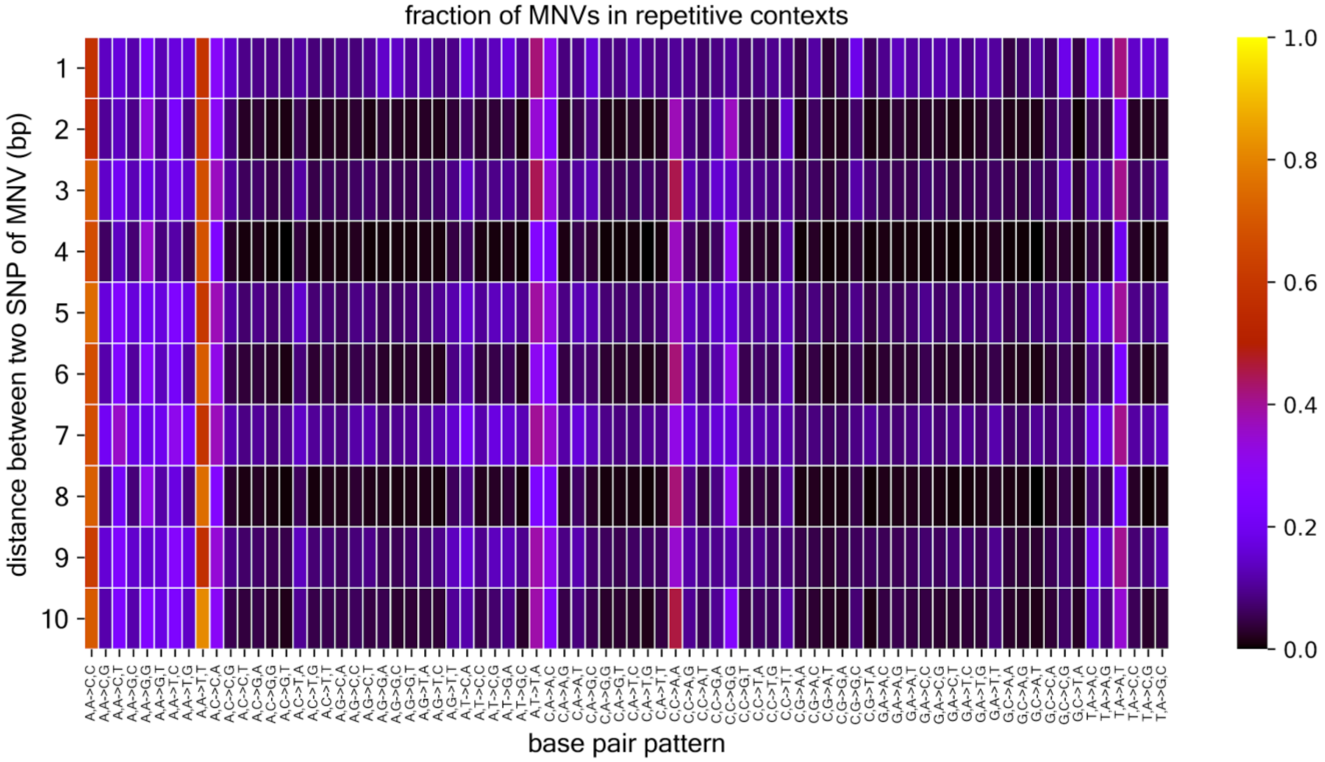


**Figure S17. Fraction of MNV that are in repetitive contexts, up to 10 bp**
Row denotes the distance between two SNPs of variant pairs, and column denotes the MNV pattern (and reverse complement). The fraction is represented as color.

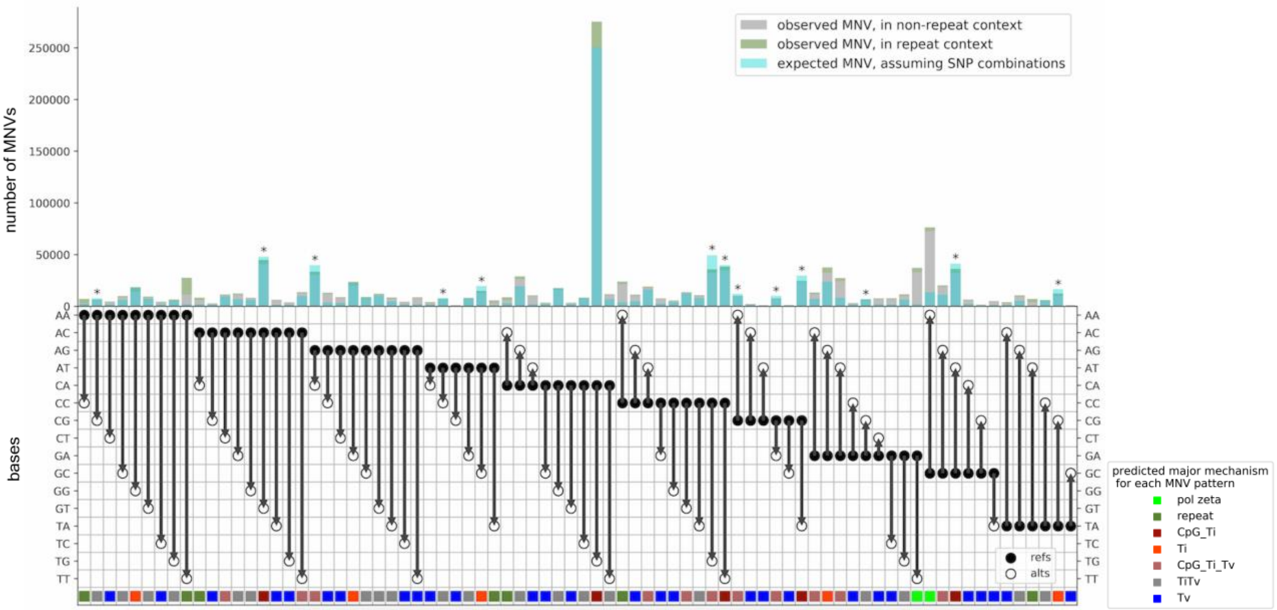


**Figure S18: Expected number of MNVs in the simulated model vs observed number of MNVs**

Cyan shows the expected number of MNVs that originates from two SNP events, when assuming that all the MNVs of pattern CA->TG in non-repeat contexts originates from two SNP events. Green is the observed number of MNVs in repeat contexts, and grey is the observed number of MNVs in non-repeat contexts. The asterisk on top of the bar indicates that the estimated number of MNVs exceeds the observed (The level of overestimation was 1.38-fold at most, for the CC->TG MNVs). The color in the bottom is as explained in figure 4.

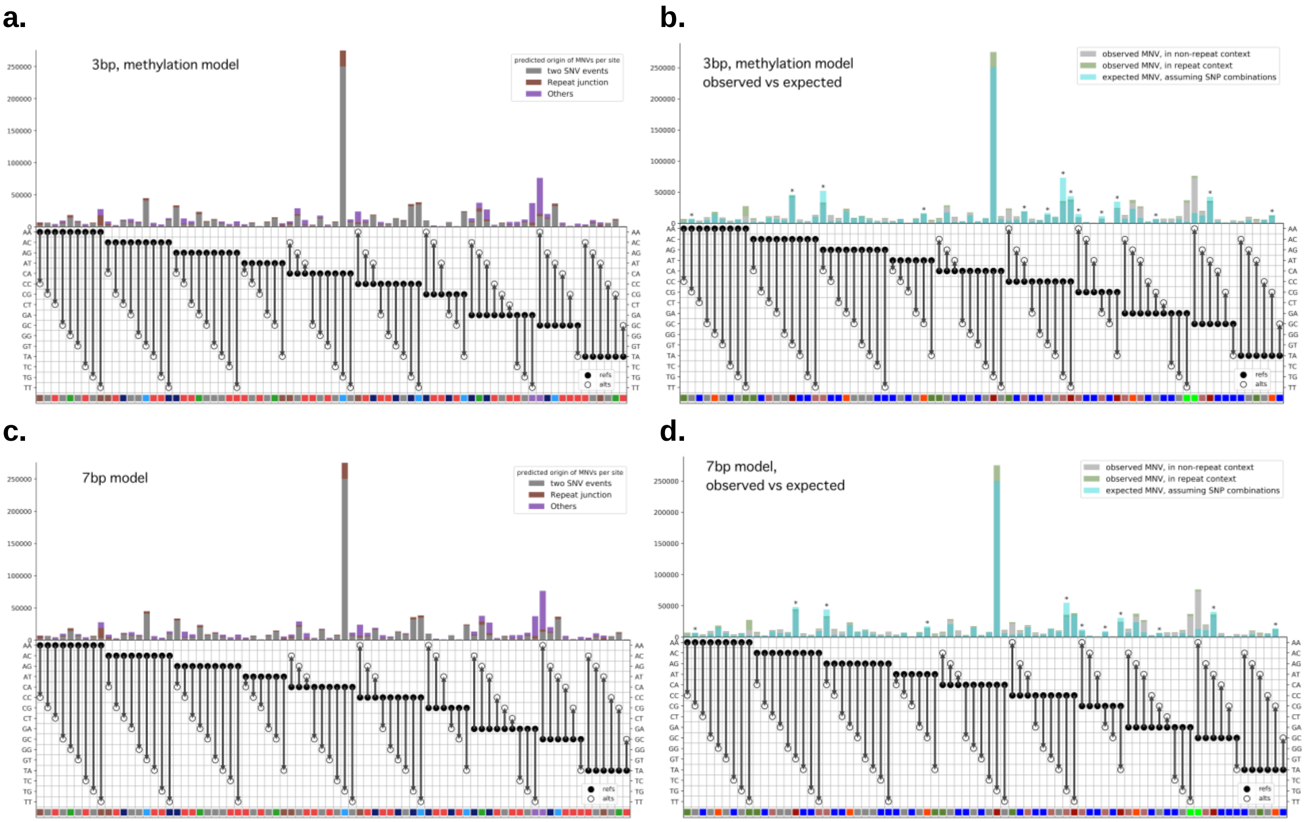


**Figure S19: MNVs in the simulated model vs observed number of MNVs, under different models**
**a,** Estimated fraction of MNVs per different biological origin, when using the methylation model. **b,** Expected number of MNVs that originates from two SNP events vs observed number of MNVs, when using methylation model. The level of overestimation for the CC->TG MNVs was 2.05-fold, higher than the canonical model. **c,** Estimated fraction of MNVs per different biological origin, when using the 7 bp context model. **d,** Expected number of MNVs that originates from two SNP events vs observed number of MNVs, when using the 7 bp context model. The level of overestimation for the CC->TG MNVs was 1.54-fold, higher than the canonical model. Axis and the color code are as explained in **figure 4** for **(a)** and **(c)**, and are as explained in the **figure S20** for **(b)** and **(d)**.

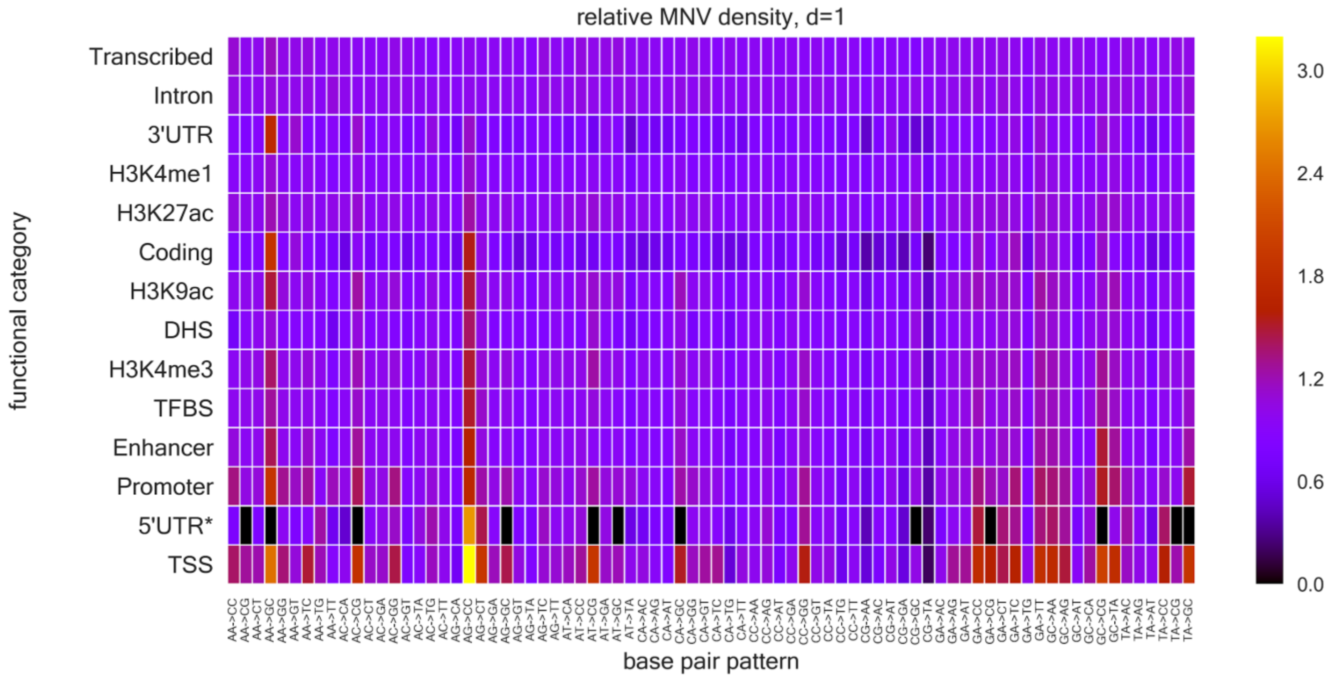
relative MNV density, d=1
Transcribed
Intron
3'UTR
H3K4me1
H3K27ac
Coding
H3K9ac
DHS
H3K4me3
TFBS
Enhancer
Promoter
5'UTR*
TSS
functional category
base pair pattern
3.0
2.4
1.8
1.2
0.6
0.0

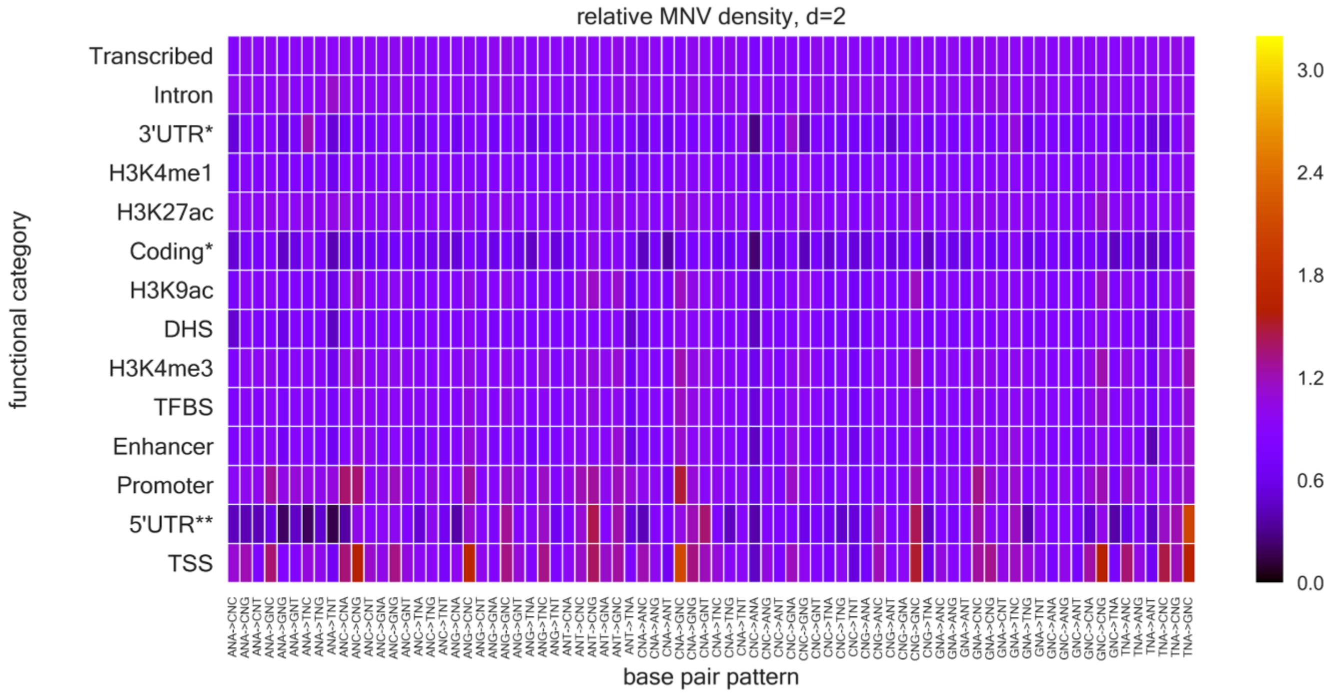
relative MNV density, d=2
Transcribed
Intron
3'UTR*
H3K4me1
H3K27ac
Coding*
H3K9ac
DHS
H3K4me3
TFBS
Enhancer
Promoter
5'UTR**
TSS
functional category
base pair pattern
3.0
2.4
1.8
1.2
0.6
0.0

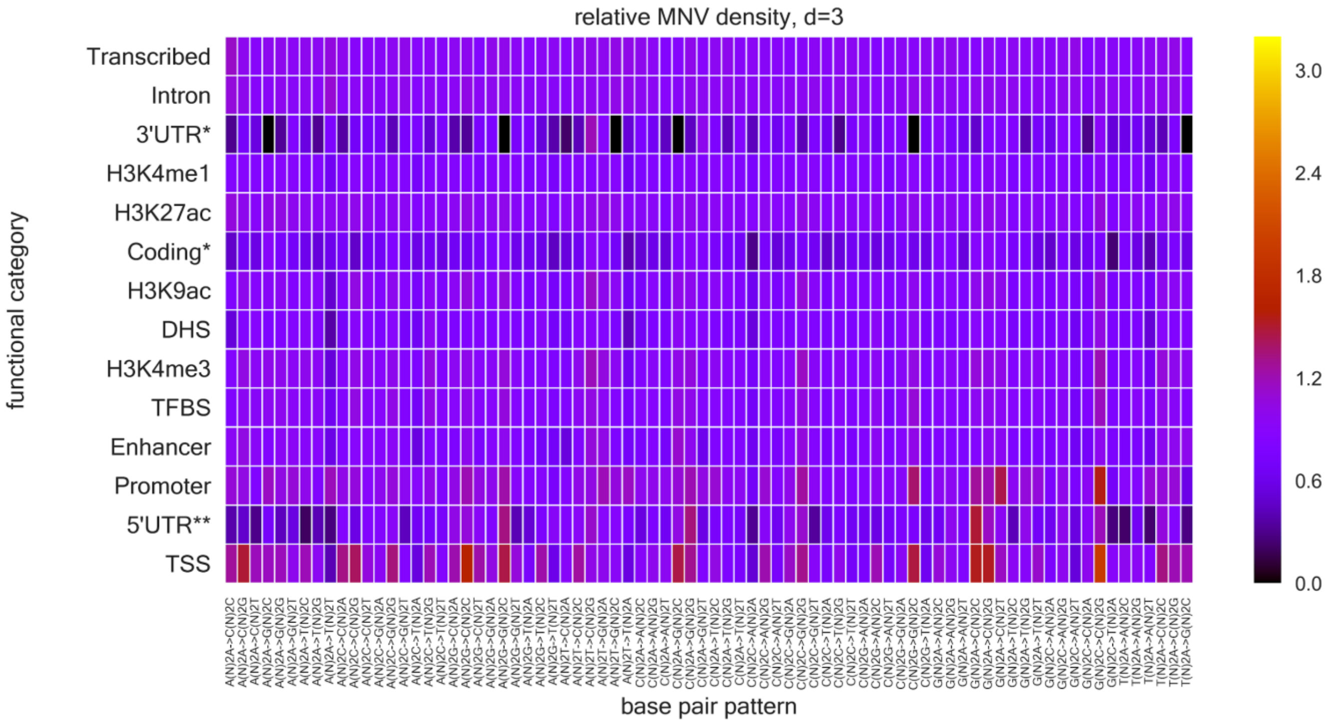
relative MNV density, d=3
functional category
Transcribed
Intron
3'UTR*
H3K4me1
H3K27ac
Coding*
H3K9ac
DHS
H3K4me3
TFBS
Enhancer
Promoter
5'UTR**
TSS
base pair pattern
3.0
2.4
1.8
1.2
0.6
0.0

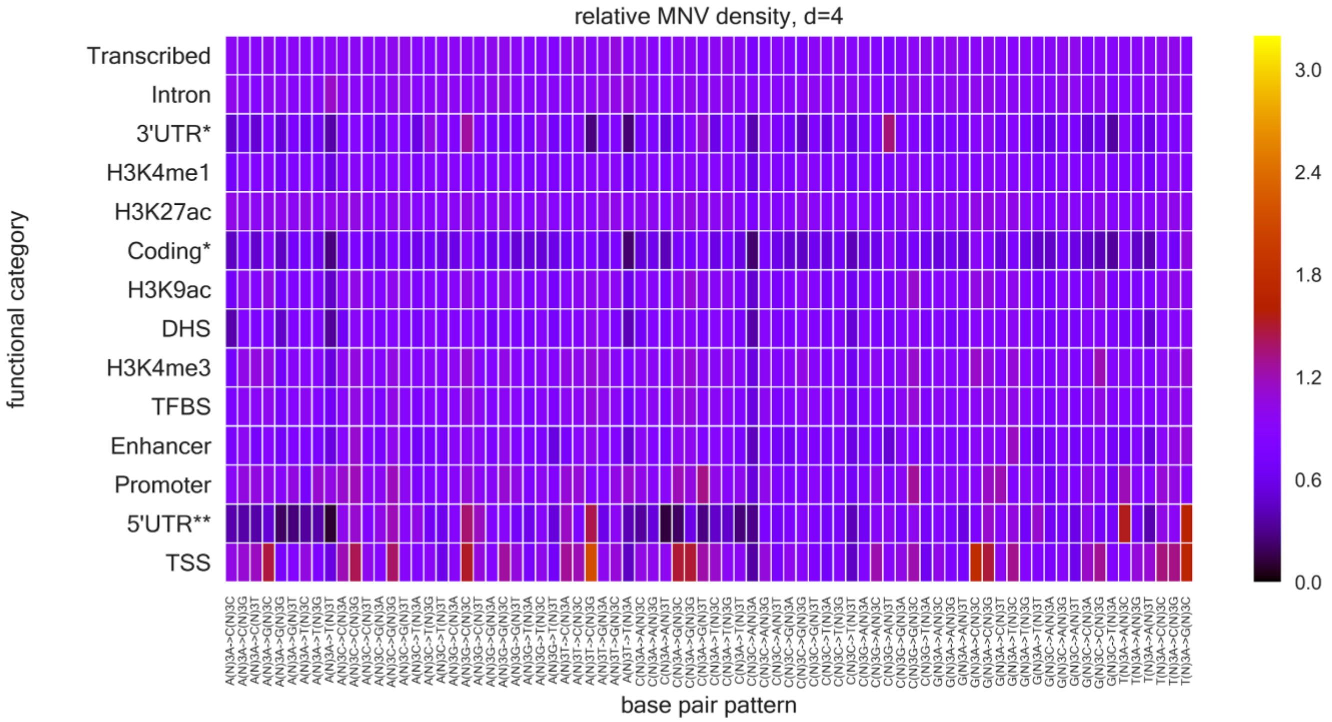
relative MNV density, d=4
functional category
Transcribed
Intron
3'UTR*
H3K4me1
H3K27ac
Coding*
H3K9ac
DHS
H3K4me3
TFBS
Enhancer
Promoter
5'UTR**
TSS
base pair pattern
3.0
2.4
1.8
1.2
0.6
0.0

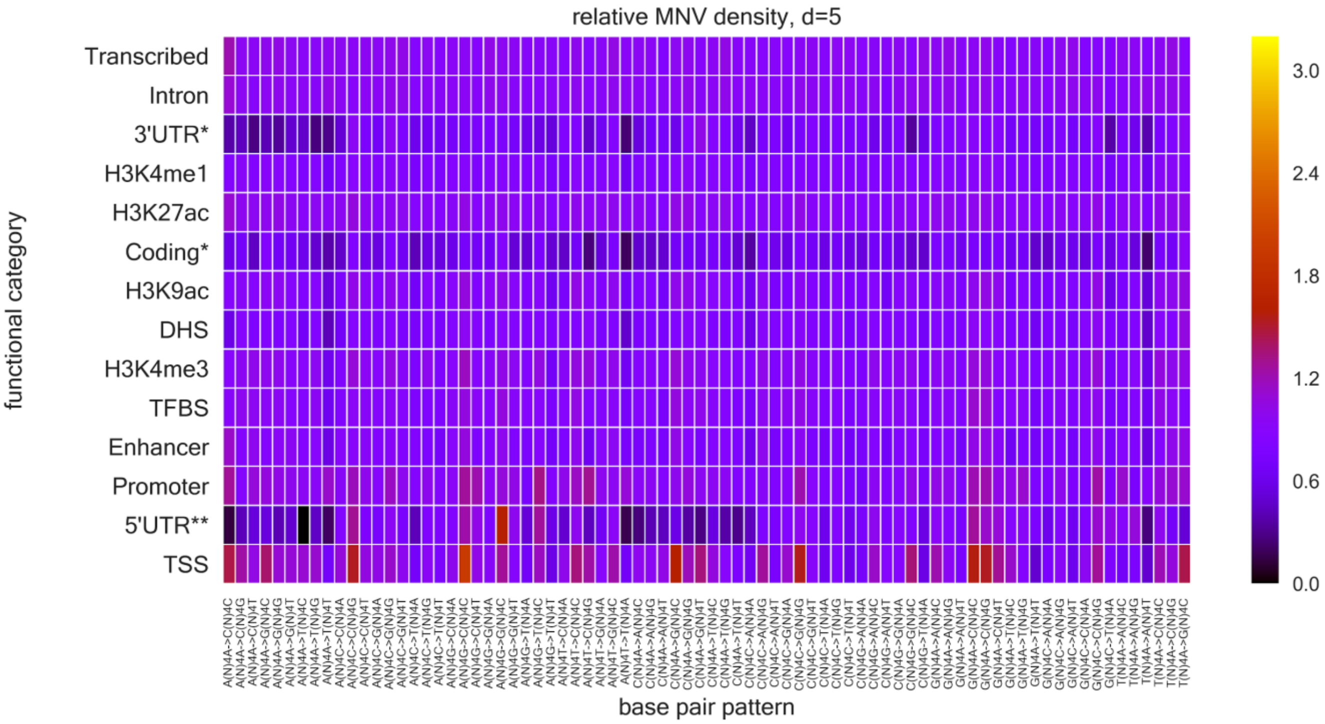
relative MNV density, d=5
Transcribed
Intron
3'UTR*
H3K4me1
H3K27ac
Coding*
H3K9ac
DHS
H3K4me3
TFBS
Enhancer
Promoter
5'UTR**
TSS
functional category
base pair pattern
3.0
2.4
1.8
1.2
0.6
0.0

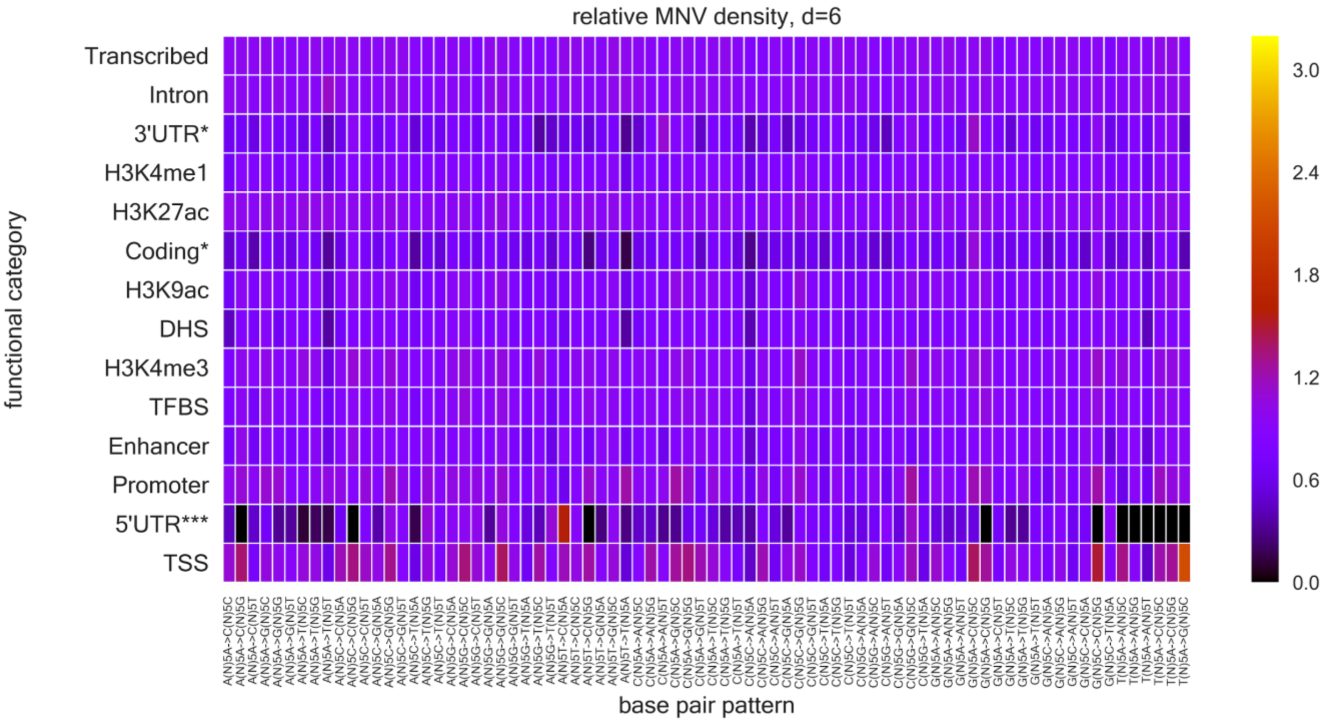
relative MNV density, d=6
Transcribed
Intron
3'UTR*
H3K4me1
H3K27ac
Coding*
H3K9ac
DHS
H3K4me3
TFBS
Enhancer
Promoter
5'UTR***
TSS
functional category
base pair pattern
3.0
2.4
1.8
1.2
0.6
0.0

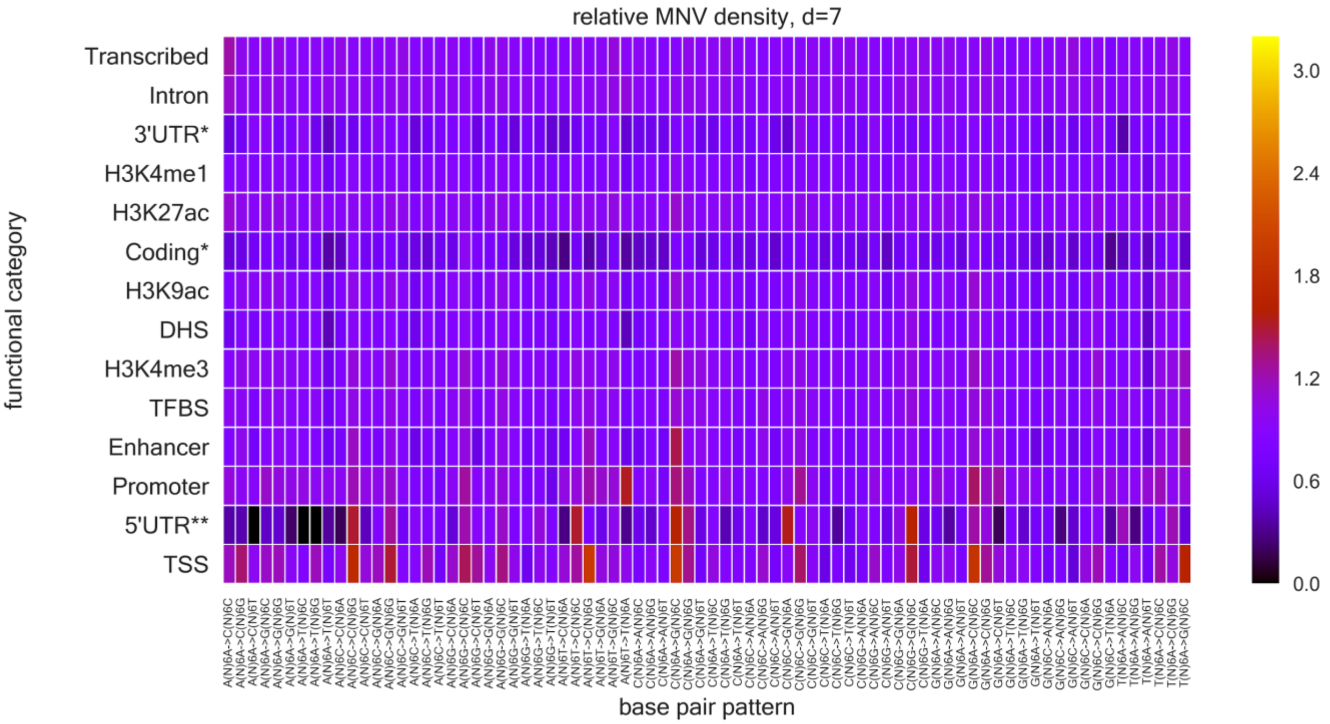
relative MNV density, d=7
functional category
Transcribed
Intron
3'UTR*
H3K4me1
H3K27ac
Coding*
H3K9ac
DHS
H3K4me3
TFBS
Enhancer
Promoter
5'UTR**
TSS
base pair pattern
3.0
2.4
1.8
1.2
0.6
0.0

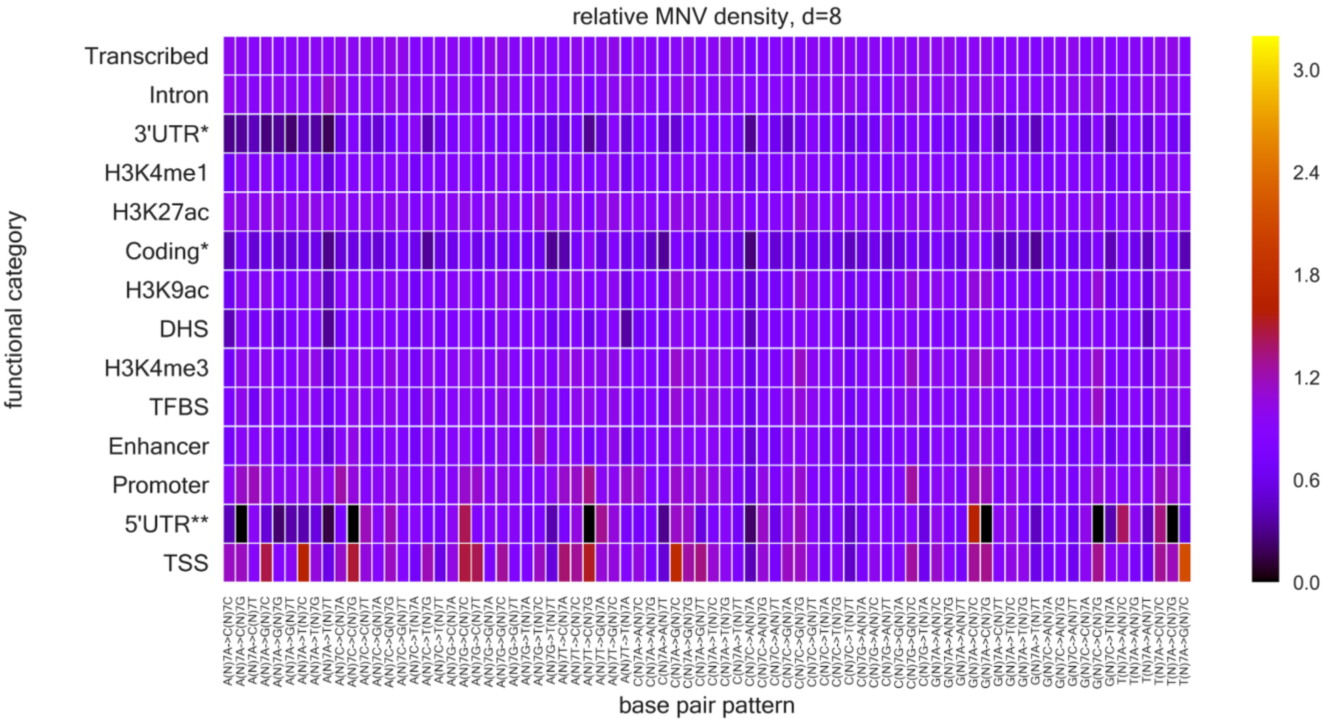
relative MNV density, d=8
functional category
Transcribed
Intron
3'UTR*
H3K4me1
H3K27ac
Coding*
H3K9ac
DHS
H3K4me3
TFBS
Enhancer
Promoter
5'UTR**
TSS
base pair pattern
3.0
2.4
1.8
1.2
0.6
0.0

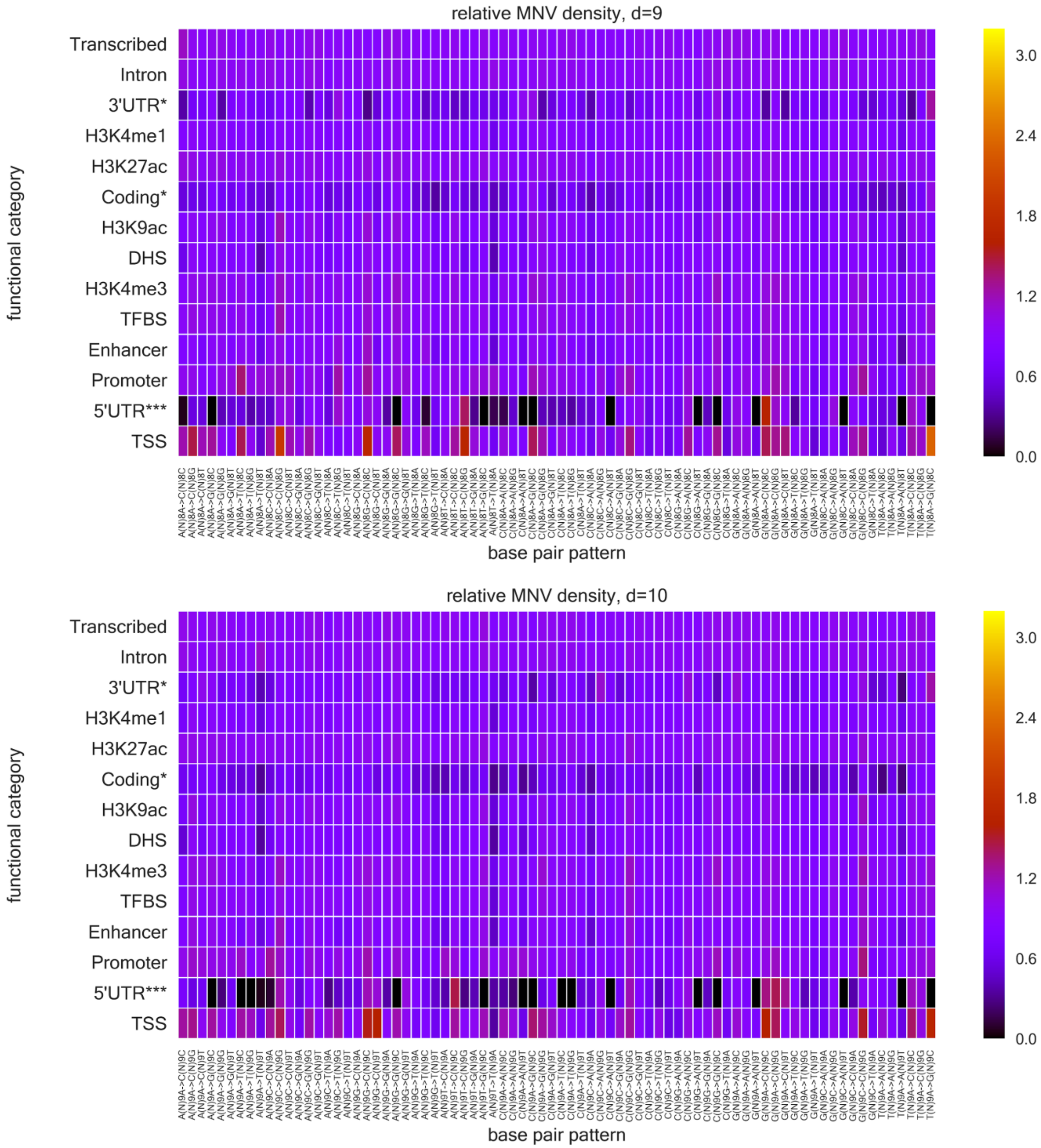


**Figure S20. Relative MNV density per functional category per MNV pattern**

Row denotes each functional category, descending order from the top by average methylation level, and column denotes the MNV pattern. The fraction is represented as color. The * in the row names shows low MNV counts (therefore limiting the statistical significance. *: <10,000 counts, **: <5,000 counts, and ***: <2,500 counts in total of 78 MNV patterns.)
